# Rapid generation of potent antibodies by autonomous hypermutation in yeast

**DOI:** 10.1101/2020.11.11.378778

**Authors:** Alon Wellner, Conor McMahon, Morgan S. A. Gilman, Jonathan R. Clements, Sarah Clark, Kianna M. Nguyen, Ming H. Ho, Jung-Eun Shin, Jared Feldman, Blake M. Hauser, Timothy M. Caradonna, Laura M. Wingler, Aaron G. Schmidt, Debora S. Marks, Jonathan Abraham, Andrew C. Kruse, Chang C. Liu

**Author notes:** These authors contributed equally to this work (Alon Wellner, Conor McMahon). Vertex Pharmaceuticals, Boston, MA 02210, USA. Department of Pharmacology and Cancer Biology, Duke University, Durham, NC 27710, USA.

## Abstract

The predominant approach for antibody generation remains animal immunization, which can yield exceptionally selective and potent antibody clones owing to the powerful evolutionary process of somatic hypermutation. However, animal immunization is inherently slow, has poor compatibility with certain antigens (*e*.*g*., integral membrane proteins), and suffers from self-tolerance and immunodominance, which limit the functional spectrum of antibodies that can be obtained. Here, we describe <u>A</u>utonomous <u>H</u>ypermutation y<u>E</u>ast surf<u>A</u>ce <u>D</u>isplay (AHEAD), a synthetic recombinant antibody generation technology that imitates somatic hypermutation inside engineered yeast. In AHEAD, antibody fragments are encoded on an error-prone orthogonal DNA replication system, resulting in *Saccharomyces cerevisiae* populations that continuously mutate surface-displayed antibody repertoires. Simple cycles of yeast culturing and enrichment for antigen binding drive the evolution of high-affinity antibody clones in a readily parallelizable process that takes as little as 2 weeks. We applied AHEAD to generate nanobodies against the SARS-CoV-2 S glycoprotein, a GPCR, and other targets. The SARS-CoV-2 nanobodies, concurrently evolved from an open-source naïve nanobody library in 8 independent experiments, reached subnanomolar affinities through the sequential fixation of multiple mutations over 3-8 AHEAD cycles that saw ∼580-fold and ∼925-fold improvements in binding affinities and pseudovirus neutralization potencies, respectively. These experiments highlight the defining speed, parallelizability, and effectiveness of AHEAD and provide a template for streamlined antibody generation at large with salient utility in rapid response to current and future viral outbreaks.

## Main Text

<u>A</u>utonomous <u>H</u>ypermutation y<u>E</u>ast surf<u>A</u>ce <u>D</u>isplay (AHEAD) pairs orthogonal DNA replication (OrthoRep) with yeast surface display (YSD) to achieve a system for the rapid evolution of antibodies. In OrthoRep, an orthogonal error-prone DNA polymerase (DNAP) replicates a special cytosolic plasmid (p1) in *Saccharomyces cerevisiae* without elevating genomic mutation rates (*1, 2*). This results in the durable continuous hypermutation of p1-encoded genes at a mutation rate of 10^−5^ substitutions per base (spb), which is 100,000-fold higher than yeast’s genomic mutation rate of 10^−10^ spb. We reasoned that when antibody fragments are encoded on p1, yeast cells would self-diversify their displayed antibodies, resulting in autonomous exploration of sequence space. When subjected to sequential rounds of sorting for antigen binding, the continuously diversifying antibodies would rapidly improve (**Fig. 1A**) and yield high-affinity, high-quality antibody clones in a short period of time. This is in contrast to typical *in vitro* antibody generation methods such as phage display (*3*) where sequence search space is defined by library complexity at the outset of an experiment but is static throughout the selection process. We first tested whether two known antibody fragments could be encoded on p1 for cell surface display. Specifically, we tested a single-chain variable fragment called 4-4-20 that binds fluorescein (*4*) and a camelid single-domain antibody fragment (“nanobody”) called AT110 that recognizes a prototypical G protein-coupled receptor (GPCR) (**Figs. 1B, 1C**) (*5*). Constitutive expression of 4-4-20 and AT110 fused to the mating adhesion receptor, Aga2p, along with induced expression of the genome-encoded cell wall anchoring subunit, Aga1p (*4*), resulted in functional display on the yeast surface as measured by flow cytometry (**Fig. 1C** and Supplementary Information). This set the stage for cycles of yeast culture and fluorescence activated cell sorting (FACS) on target binding to effect the rapid evolution of high-affinity antibodies, akin to somatic hypermutation for affinity maturation in vertebrate immune systems (*6, 7*).

**Fig 1.**
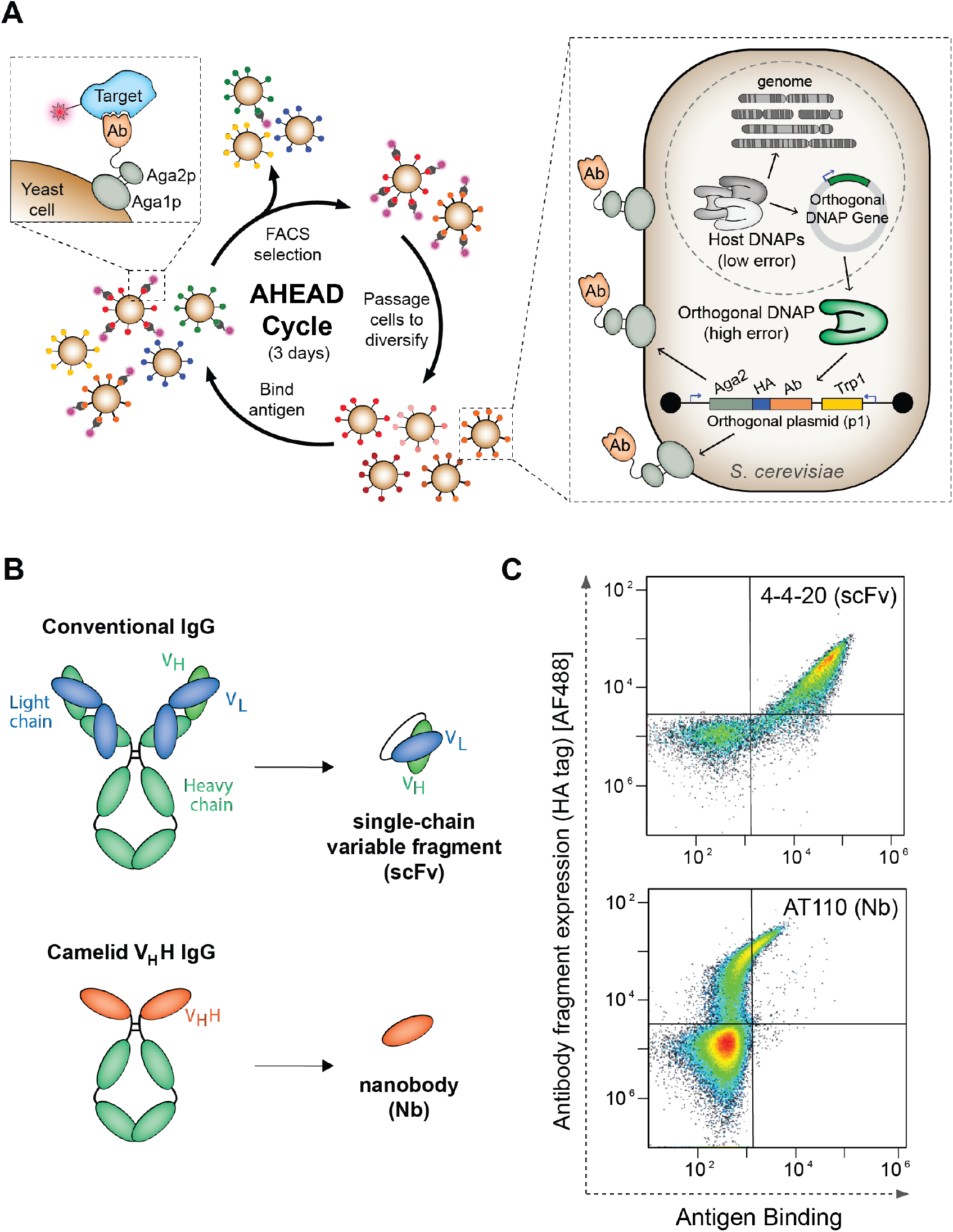
<u>A</u>utonomous <u>H</u>ypermutation y<u>E</u>ast surf<u>A</u>ce <u>D</u>isplay (AHEAD). (**A**) Scheme for rapid evolution of high-affinity binding using AHEAD. Ab = antibody, DNAP = DNA polymerase, HA = hemagglutinin tag. (**B**) Antibody fragments used in this study and their relationship to conventional antibodies. (**C**) Cytometry plot showing detection of a functionally surface-displayed scFv and a functionally surface-displayed Nb encoded on the p1 orthogonal plasmid, replicated by an associated orthogonal DNAP. The orthogonal DNAP used in this case was the wt TP-DNAP1 (see Supplementary Information) rather than the error-prone TP-DNAP1-4-2 variant that was used for all subsequent AHEAD evolution experiments. Cognate antigens for 4-4-20 (fluorescein) and AT110 (AT1R) were labeled with biotin and FLAG tag, respectively, and detected with APC-conjugated anti-FLAG or AF647-conjugated streptavidin, respectively. The HA tag was detected with mouse anti-HA and a goat anti-mouse AF488-conjugated secondary antibody.

To test whether AHEAD could generate high-affinity antibodies through cycles of yeast culture and FACS, we sought to improve the nanobody, AT110. Nanobodies are single-domain antibody fragments derived from the V_H_H domains of heavy chain antibodies in llamas, camels, and their relatives (**Fig. 1B**) (*8*). These unique antibody fragments are capable of stabilizing distinct functional conformations of GPCRs and other dynamic proteins by virtue of their propensity to bind concave epitopes (*9*). This has made the discovery of high-affinity nanobodies a mainstay of GPCR structural biology and pharmacology (*10, 11*). Nanobodies are also particularly appropriate for animal-free antibody generation approaches, as the alternative of immunizing camelids carries logistical challenges of large animal husbandry and animal welfare considerations (*12*). AT110 is a low-affinity nanobody which binds selectively to the active-state conformation of the angiotensin II receptor type 1 (AT1R), discovered *via in vitro* selection from a naïve synthetic nanobody library. However, AT110 required several rounds of manual error-prone PCR diversification, selection, and engineering to reach the affinity necessary for co-crystallization and structural studies (*5, 13*). Using AHEAD and starting from AT110, we carried out iterative cycles of yeast culture and FACS to improve AT110 (**Fig. 2A**). This experiment yielded nanobody AT110i104 (**Table S1**), with an allosteric modulation potency of 2.5 nM for enhancing agonist binding to the AT1R GPCR (**Fig. 2B**), representing a roughly 20-fold functional affinity enhancement compared to the parent clone. Notably, some mutations (*e*.*g*., I98V and a non-synonymous change at Y113) were identified both by AHEAD and previous efforts (*5*), while other affinity-enhancing mutations (*e*.*g*., R45C and R66H) were distinct (**Fig. S1**). Interestingly, Y113H and I98V were synergistic in their ability to modulate agonist-bound AT1R (**Fig. 2B**), showing that AHEAD could find complex functional outcomes. Overall, this experiment demonstrated that AHEAD can produce high-affinity antibodies in a much more streamlined manner than conventional approaches.

**Fig 2.**
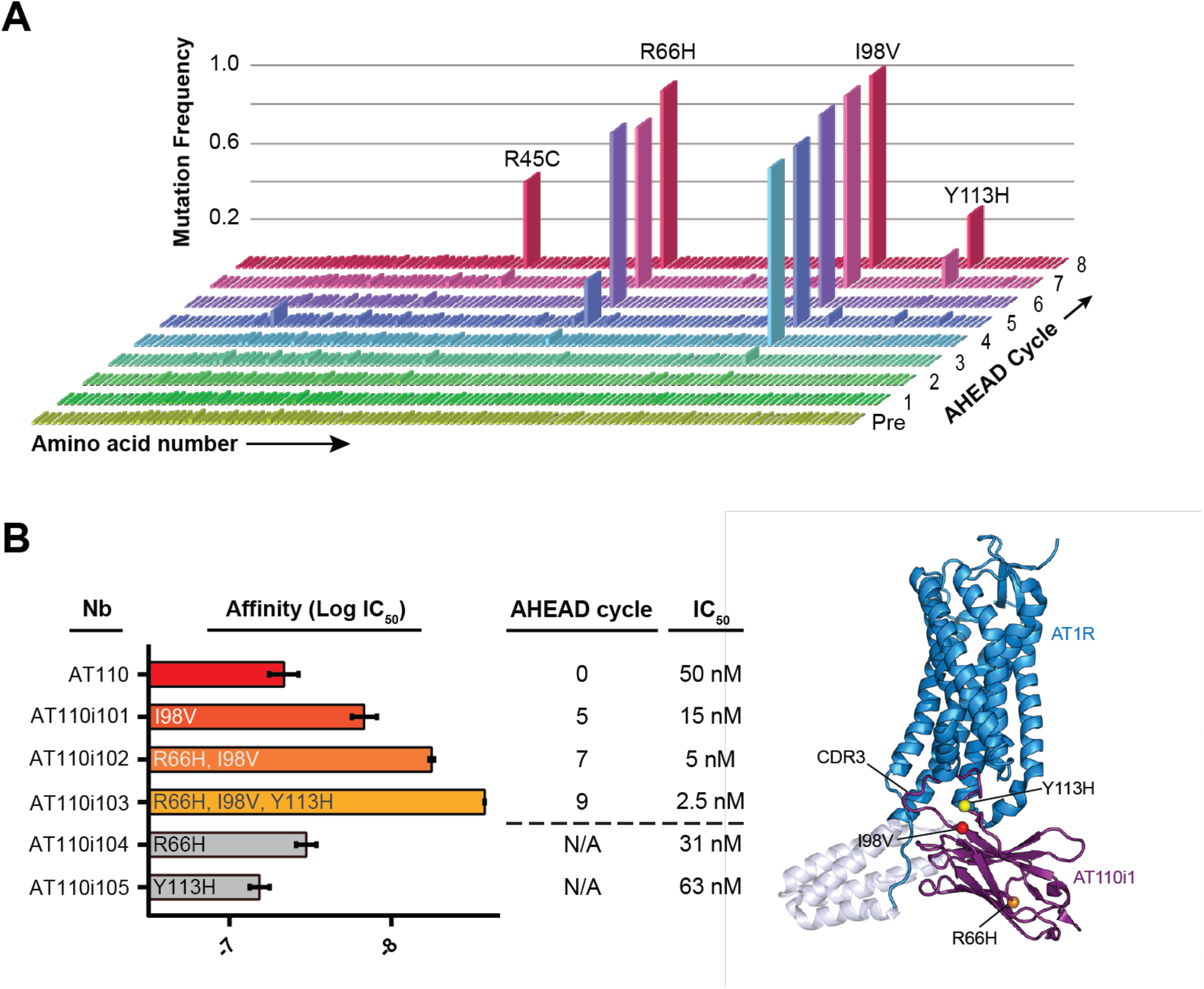
Evolution of anti-AT1R nanobodies. (**A**) Enrichment of affinity-increasing mutations in anti-AT1R nanobodies through cycles of AHEAD as determined by NGS of the p1-encoded nanobody population in each cycle of AHEAD. Pre = population composition before the first cycle of AHEAD. (**B**) Nanobody potency was assessed in a radioligand allosteric shift assay. This measures the ability of each nanobody to enhance agonist affinity by stabilizing an active-state receptor conformation, serving as an indirect measure of nanobody binding affinity. Error bars represent the SEM from three independent experiments performed as single replicates.

Although successful, lessons learned during anti-AT1R nanobody evolution motivated us to redesign AHEAD’s nanobody display constructs (**Fig. S2A**). In particular, we found that the average level of AT110 display was low, which complicated the FACS selection process in cycles of AHEAD. We overcame this problem by using magnetic activated cell sorting (MACS) before each FACS round in order to enrich subpopulations of cells with high AT110 display. Since OrthoRep maintains p1 in multiple copies (*1*), copy number fluctuations create subpopulations that express substantially more nanobody than average, allowing effective MACS enrichment of these higher expressers when large numbers of cells are used. However, the requirement for MACS steps significantly increased the time and effort needed for antibody evolution, counter to AHEAD’s potential as a rapid and streamlined antibody generation system. To obtain a general solution to suboptimal nanobody display, we adopted an improved display architecture (*14*), evolved a new p1-specific promoter, engineered a stronger secretory leader, and utilized a polyadenosine tail (*15*) encoded downstream of the nanobody gene for AHEAD (**Figs. S2A-C** and Supplementary Information). These features together increased the expression and display level of nanobodies by ∼25-fold, allowing the typical cell in a population to display enough nanobody for direct antigen binding selection by FACS even when binding was weak (**Fig. S2D**). This second-generation AHEAD system supports our intended process of cycling directly between yeast culture and FACS to generate antibodies (**Fig. 1A**).

To test our upgraded system, we ran several antibody fragment evolution experiments against additional targets including human serum albumin (HSA) and green fluorescent protein (GFP). High-affinity nanobodies against HSA are desired as domains that can be fused to therapeutic proteins in order to increase their serum half-life (*16*). High-affinity nanobodies against GFP are useful for immunostaining in GFP-expressing tissue samples where conventional antibodies take too long to penetrate due to their high molecular weight (*i*.*e*. cleared brain) (*17, 18*). Starting from relatively low-affinity nanobodies targeting HSA and GFP, namely Nb.b201 (*19*) and Lag42 (*20*), respectively, we successfully evolved high-affinity clones through 4-6 cycles of AHEAD (**Fig. S3** and **Table S1**).

AHEAD is distinguished by its coincident speed, parallelizability, and evolutionary process for antibody maturation. These properties are particularly valuable for outbreak response where urgency demands rapid discovery (speed) of high-affinity antibodies through many independent antibody generation campaigns that collectively maximize the probability of success (parallelizability). In light of the current COVID-19 pandemic, we asked whether AHEAD would be capable of generating collections of potent nanobodies against the novel coronavirus, SARS-CoV-2 (*21*). Starting from an open-source naïve nanobody YSD library (*19*), we selected 8 clones that bound the receptor-binding domain (RBD) of the SARS-CoV-2 spike (S) protein (**Fig. S4A** and **Table S1**). Each of these distinct clones was then encoded on p1 for streamlined evolution (*i*.*e*., affinity maturation). The parallelizability of AHEAD allowed us to simultaneously run 8 independent affinity maturation experiments, one for each of the parental clones, in order to avoid clonal interference among initial lineages and increase the chance of obtaining matured nanobodies with functional diversity in binding modes (**Fig. S4B**) (*22*). After 3-8 cycles of AHEAD (**Fig. S5**) where the uninterrupted cycle time defined by yeast culturing and FACS was only 3 days, all eight experiments produced multi-mutation nanobodies (**Figs. 3A, 3B** and **Table S1**) with higher affinity for RBD than their parents (**Fig. 3C, Table 1, Table S2** and **Fig. S6**). For example, RBD1i13, RBD3i17, RBD6id, RBD10i10, RBD10i14, and RBD11i12 exhibited monovalent RBD-binding affinity improvements up to ∼580-fold over the course of affinity maturation (**Fig. 3** and **Table 1**), and one nanobody, RBD10i14, reached a subnanomolar monovalent *K*_d_ of 0.72 nM (**Fig. 3C** and **Table 1**). These levels of evolutionary improvement from naïve nanobodies, achieved through a fast and parallelizable process, confirm the power of AHEAD.

**Table 1.** Performance of anti-SARS-CoV-2 nanobodies (Nbs) evolved using AHEAD. See **Table S2** for complete information.

| Nb Name | AHEAD cycle | Mutations | $K_d$ (nM) | Neutralization $IC_{50}$ ($\mu g\ mL^{-1}$) | ACE2 competition |
| --- | --- | --- | --- | --- | --- |
| RBD1 | 0 | wt | 1400 | 3.76 | N.D.<sup>1</sup> |
| RBD1i1 | 4 | E46K, T100A | 48.1 | 0.18 | Strong |
| RBD1i13 | 7 | E44G, E46K, K86E, T100A | 32.2 | 0.05 | Strong |
| RBD3 | 0 | wt | 6000 | >26 | N.D. |
| RBD3i2 | 4 | V2A, Y58R | 128 | 5.52 | N.D. |
| RBD3i17 | 8 | V2A, Y58R, Y59C | 230 | 0.116 | N.D. |
| RBD6 | 0 | wt | 990 | 0.25 | N.D. |
| RBD6id | 3 | S21N, S25N, D61I | 263 | 0.056 | Moderate |
| RBD10 | 0 | wt | 417 | >45 | N.D. |
| RBD10i10 | 4 | E44G, E46K, S55G, D61G | 2.14 | 0.19 | None |
| RBD10i14 | 7 | E44G, E46K, M34V, D61G | 0.72 | 0.42 | N.D. |
| RBD11 | 0 | wt | 2420 | >37 | N.D. |
| RBD11i12 | 4 | E44G, H105Y, G109S | 316 | 0.04 | Moderate |

<sup>1</sup>N.D. = Not Determined

**Fig 3.**
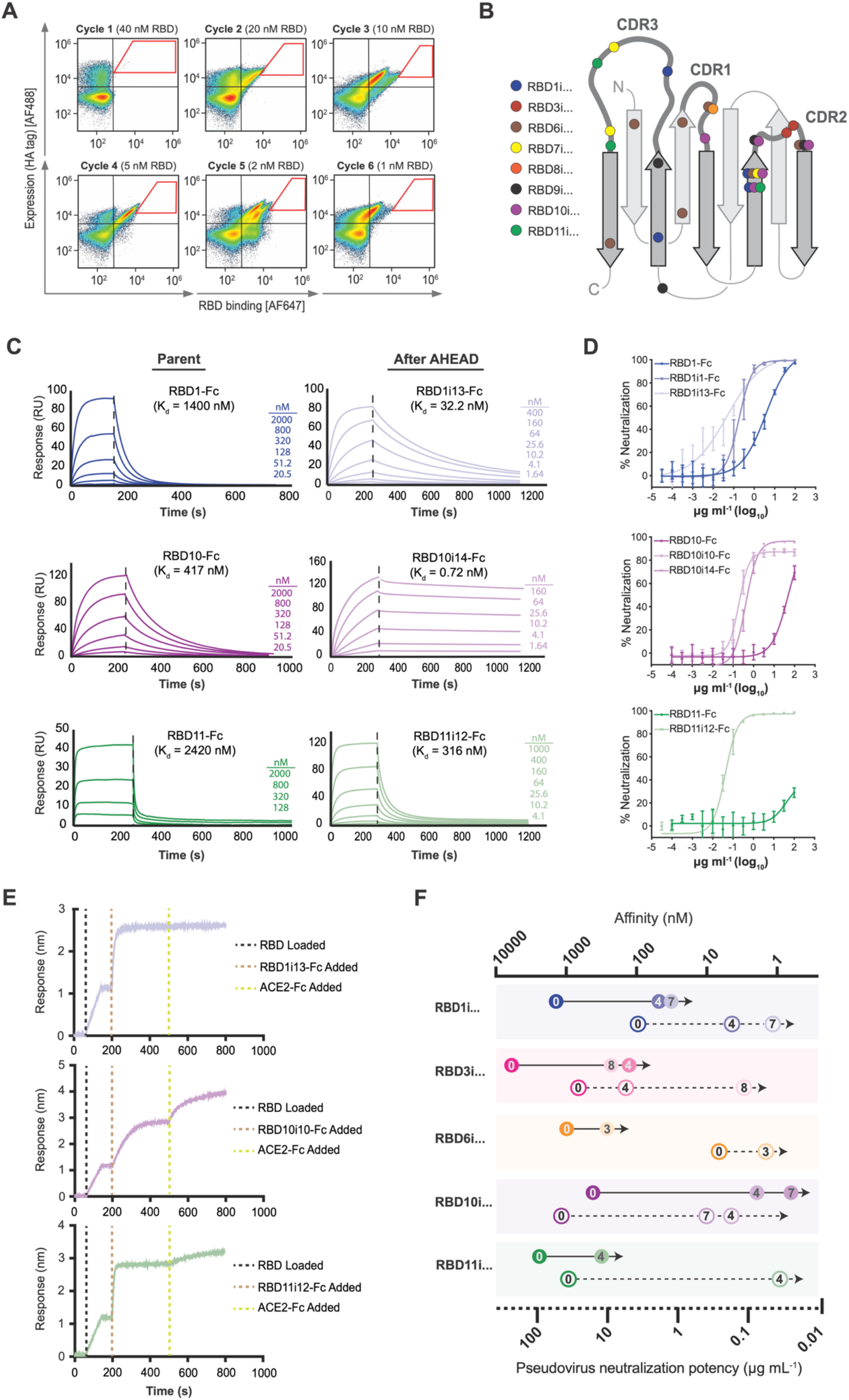
Evolution of anti-SARS-CoV-2 nanobodies and activities of highly potent anti-SARS-CoV-2 nanobodies. (**A**) Representative FACS plots showing affinity maturation of an anti-SARS-CoV-2 nanobody (Parent = RBD10). Red polygons correspond to gates used for sorting. (See **Fig. S5** for similar plots showing affinity maturation from all parents.) (**B**) Location of nanobody mutations fixed in 8 independent AHEAD experiments starting from different parental clones. (**C**) Surface plasmon resonance (SPR) traces and associated monovalent affinities of select anti-SARS-CoV-2 nanobodies evolved using AHEAD. (See **Fig. S6** for affinity measurements on additional nanobodies.) (**D**) Neutralization activities of select anti-SARS-CoV-2 nanobodies on pseudotyped SARS-CoV-2 virus. Each nanobody concentration (X-axis) was tested in replicate. N = 6, error bars represent ± s.d. (See **Fig. S7A** for neutralization activities of additional nanobodies.) (**E**) Bio-layer interferometry (BLI) traces measuring ACE2 competition for RBD binding in the presence of select anti-SARS-CoV-2 nanobodies evolved using AHEAD. (See **Fig. S7B** for ACE2 competition activities of additional nanobodies and control antibodies.) (**F**) Affinity and neutralization potency improvements of nanobodies isolated from different cycles of AHEAD during each parent nanobody’s affinity maturation. Each closed circle represents a nanobody’s affinity and the open circle of identical color represents the nanobody’s neutralization potency. The number within each circle designates the AHEAD cycle from which the nanobody was isolated.

Anti-RBD monoclonal antibodies have emerged as promising therapeutics for the treatment of COVID-19 (*23, 24*). These antibodies act by inhibiting the interaction between SARS-CoV-2 and its receptor, angiotensin-converting enzyme 2 (ACE2), thereby blocking entry into cells. To probe the potential of our anti-RBD nanobodies as therapeutic candidates, we genetically fused the nanobodies to the Fc region of the human IgG1 antibody isotype and carried out neutralization assays against SARS-CoV-2-pseudotyped virus. We found that many of the evolved anti-RBD nanobodies had exceptional neutralization potencies representing up to ∼925-fold improvements over their parent sequences (**Fig. 3D, Table 1**, and **Fig. S7A**). For example, RBD1i13, RBD3i17, RBD6id, RBD10i10, RBD10i14, and RBD11i12 exhibited low nanomolar or subnanomolar half-maximal inhibitory concentration (IC_50_) values of 0.66, 1.51, 0.72, 2.44, 5.38, and 0.52 nM, respectively (**Fig. 3D, Table 1, Table S2**, and **Fig. S7A**). Interestingly, RBD1i13 and RBD11i12, which had the strongest viral neutralization potencies among all evolved variants, descended from parents that were relatively poor neutralizers (**Table 1** and **Table S2**). This highlights the value of experimental parallelizability available to AHEAD: through evolution experiments that kept each parent clone’s lineages separate, early high achievers such as RBD6 could not outcompete the initially low performing lineages that ultimately gave rise to the most potent neutralizers. This is akin to ‘demes’ in natural evolution, which act to reduce clonal interference and increase overall functional diversity (*25, 26*) Indeed, AHEAD’s ability to maximize diversity of high-affinity clones through independent experiments should be valuable in all antibody generation campaigns, given that antibody performance depends on secondary features beyond affinity alone.

To assess the functional diversity of nanobody clones from AHEAD, we tested potent neutralizers for their ability to compete with ACE2 binding. Clones RBD1i13, RBD6id, and RBD11i12 strongly or moderately competed with ACE2 binding whereas a fourth clone, RBD10i10, did not compete at all (**Fig. 3E, Table 1**, and **Fig. S7B**). This suggests that the sites of RBD binding and mechanisms of neutralization differ across our collection of potent nanobodies, which could be exploited in nanobody cocktails or fusions between different nanobodies. Such strategies can lead to higher neutralization potency or limit mutational escape as SARS-CoV-2 evolves in the field (*27*).

The rapid evolution of anti-SARS-CoV-2 nanobodies where 8 independent antibody generation experiments were carried out in an uninterrupted time of only 1.5-3 weeks that saw major improvements in both affinity and neutralization potencies (**Fig. 3F** and **Table S2**) may serve as a template for future outbreak response. Since we started from naïve synthetic nanobody libraries, our experiments did not depend on the prior discovery of antibodies from patients or animals, demonstrating capacity for immediate response once a molecular target is identified. Since we ran multiple independent evolution campaigns simultaneously, we readily obtained a collection of functionally diverse nanobody sequences, important for hedging against biological uncertainty in the face of new pathogens going forward. Since we relied only on simple cycles of yeast culturing and FACS to evolve potent nanobodies, our experiments should be broadly accessible, and may enable wider and more distributed antibody development efforts in future pandemics. In the particular case of anticipating novel coronavirus outbreaks, our collection of SARS-CoV-2 nanobodies, already encoded on AHEAD, should be a privileged starting point for rapid response as additional evolutionary cycles may be able to direct our current nanobodies to become specific for new spike variants. In fact, we estimate that given an SARS-CoV-2 RBD escape variant (*28*) we would be able to discover multiple high-affinity binders in 2 weeks.

Beyond outbreak response, AHEAD has the potential to broadly upgrade antibody generation supporting all areas of biomedicine, especially as the AHEAD technology matures. For example, we foresee expansions of AHEAD where instead of starting evolution from isolated initial clones encoded on p1, diverse naïve antibody collections are encoded as libraries. Such off-the-shelf libraries of yeast would allow AHEAD experiments to encompass both initial clone discovery and affinity maturation in an end-to-end process for antibody generation against arbitrary targets. To test the feasibility of this possibility, we carried out experiments where a 200,000-member naïve nanobody library capturing key features of camelid immune repertoires was computationally designed and synthesized (*29*), encoded in AHEAD strains with 50-fold coverage, and subjected to selection for binding GFP as a test target. After three cycles of AHEAD, a single nanobody, NbG1, dominated the population, and after two additional cycles, a C96Y mutation that increased GFP binding (EC_50_) by 4.4-fold arose and fixed (**Fig. S8** and **Table S2**), emulating the entire process of somatic recombination, clonal expansion, and somatic hypermutation in the immune system. As more libraries and developments in the underlying OrthoRep system (*e*.*g*., higher mutation rate orthogonal DNAPs) become available, AHEAD’s core functionality as a rapid and parallelizable affinity maturation process involving only the culturing and sorting of yeast will form a new foundation for antibody generation.

## Materials and Methods

### AHEAD base yeast display strain construction

We generated strain yAW301, which is essentially the EBY100 yeast surface display strain (*4*) harboring a ‘landing pad’ version of p1 along with p2 (*30*). yAW301 served as the base strain for all AHEAD experiments where 1) antibody fragment display cassettes were integrated onto the landing pad p1 and 2) a nuclear plasmid, pAR-633-Leu2 (**Table S3**), encoding an error-prone orthogonal DNAP replicating p1, was transformed into the strain in order to drive continuous hypermutation of p1 and the antibody fragment display cassette encoded therein. To make strain yAW301, we first generated F102-2 (*30*) cells containing p1 and p2 but lacking the MET17 gene from the genome. To delete MET17, F102-2 cells were transformed with a linear DNA fragment encoding a G418 resistance cassette flanked on both sides by 45bp sequences homologous to the surrounding regions of MET17 (SG ID S000004294). Following selection on solid media with G418, colonies were isolated and verified for the deletion of MET17 by PCR. In addition, the colonies’ inability to grow without the supplementation of methionine and cysteine was verified. F102-2 cells harbor the unmodified cytosolic p1 and p2 plasmids. However, we wished to engineer a version of p1 designed for ease of integration and selection of antibody yeast display cassettes. To generate such a ‘landing pad’ p1, a DNA fragment was designed to recombine with F102-2’s p1 plasmid (8.9kb) in order to replace positions 3201-8400 of p1 with MET17 driven by the p1 specific promoter, p2O5. This integration resulted in the desired landing pad p1 (5.3kb) harbored in F102-2. The shifted size of p1 was validated by gel electrophoresis of total DNA treated with proteinase K (Thermo Fisher, USA), as previously described (*30*), and sequenced verified. This landing pad p1 could then be subsequently transported from F102-2 to other strains through protoplast fusion, as previously described (*30*). In particular, EBY100 yeast surface display cells (*4*) that had their genomic MET17 deleted using the same strategy for deleting MET17 in F102-2 were fused with F102-2 cells containing the landing pad p1. Selection for prototrophies uniquely encoded by nuclear genes in EBY100 (and not in F102-2) as well as selection for MET17 uniquely encoded on p1 resulted in strain yAW292 (**Table S4**). Finally, plasmid pAR-633-Leu2, which encodes the error-prone orthogonal DNAP, TP-DNAP1-4-2 (*2*), was transformed into yAW292 so that the p1-encoded nanobody expression cassette would be replicated by the error-prone orthogonal DNAP to drive hypermutation. The resulting strain was dubbed yAW301 (**Table S4**).

### Cloning nanobodies into AHEAD

pAW24 or pAW240, plasmids that encode the linear cassette needed for integration of nanobodies into the landing pad p1, were designed and constructed for nanobody expression from p1 (**Fig. S2** and **Table S3**). In pAW24 (corresponding to AHEAD 1.0, see **Fig. S2**), the nanobody gene within the integration cassette is driven by a previously reported p1-specific promoter called p10B2 (*15*) and fused to a standard alpha mating factor secretory leader sequence (*4*). In pAW240 (corresponding to AHEAD 2.0, see **Fig. S2**), the nanobody gene within the integration cassette is driven by a new p1-specific promoter (pGA) that was discovered in an unrelated OrthoRep-based protein evolution project. (pGA is the same sequence as p10B2 with two G->A mutations at positions −5 and −34 upstream from the start codon.) The nanobody gene within the integration cassette of pAW240 also includes a mutated leader sequence, app8i1, which was selected for higher efficiency display (**Fig. S2C**). Finally, the nanobody expression cassette in pAW240 contains a hard-coded poly-A tail (*15*) to maximize expression. Single nanobodies or libraries were amplified with PCR using primers Nb_P240_F and Nb_P240_R (**Table S5**), gel purified and assembled into BseR1-digested pAW24 or pAW240 with a Gibson reaction. For pAW240, it is important to use BseR1 digested pAW240 as the “backbone” for Gibson assembly as opposed to a PCR amplified “backbone” of pAW240 because PCR causes truncation of the hard-coded poly-A tail sequence, which results in lower expression and a drop in antibody display levels. Once pAW24 or pAW240 were properly assembled to contain desired nanobodies, the resulting plasmids were linearized with ScaI to expose flanking regions homologous to the landing pad p1 to direct their homologous recombination into p1 in yAW292 or yAW301. The ScaI linearized pAW24 or pAW40 plasmids containing desired nanobodies were then transformed into yAW301 by a standard yeast chemical transformation protocol (*31*). Cells that successfully integrated the nanobody expression cassette onto p1 were selected on Synthetic Complete without Histidine, Leucine, Uracil, Trptophan, Methionine, and Cysteine (US Biological) with 2% glucose (SC-HLUWMC) agar plates. Although MET17 in the landing pad p1 is replaced by the nanobody expression cassette, we found that the exclusion of methionine was appropriate for selection, since some copies of the landing pad p1 would still be present in cells containing the p1 integration product. We empirically found that selection for cells still containing the unmodified landing pad p1 increased the success rate of integration, possibly because the unmodified landing pad p1 also encodes an additional source of TP-DNAP1, allowing for a higher overall p1 copy number to aid the replication of both the unmodified landing pad p1 and the modified nanobody-expressing p1 integration product. Proper integration of the nanobody expression cassette into the landing pad p1 was varied by gel electrophoresis on total DNA, PCR using primers unique to the integration product, and DNA sequencing. For further cultivation, cells were grown in SC-HLUW (Synthetic Complete without Histidine, Leucine, Uracil, and Trptophan with 2% glucose, US Biological) liquid dropout media. AHEAD experiments were started after this point.

### Affinity maturation of anti-AT1R nanobodies using the first-generation AHEAD 1.0 system

Yeast strain yAW301 harboring pAR-633-Leu2 encoding the error-prone orthogonal DNAP and an AT110 expression cassette (linearized plasmid pAW24) integrated onto p1 were grown and passaged in SC-HLUW. In preparation for each fluorescence-activated cell sorting (FACS) selection step, display of AT110 was induced by transferring yeast to SC-HLUW but where glucose was replaced with 2% galactose as the sole sugar source. This results in the induction of Aga1p expression and subsequent surface display of the AT110 nanobody. Because of the low expression of nanobodies in the first-generation AHEAD 1.0 system (**Fig. S2**), cells expressing the highest levels of AT110 nanobodies were deliberately enriched before each AHEAD cycle. Accordingly, the following process defined each AHEAD 1.0 cycle, which is longer than the 3-day AHEAD 2.0 cycle used for the majority of evolution experiments in this work. Between 2 x 10^9^ and 1 x 10^10^ induced yeast were pelleted and resuspended in 2.5-5 mL of AT1R staining buffer (20 mM HEPES pH 7.5, 150 mM NaCl, 0.1% BSA, 3 mM CaCl2, 0.1% MNG, 0.01% CHS, 0.2% maltose, 20 µM angiotensin II) with 1 µM anti-HA antibody that was fluorescently labeled, alternately, with AlexaFluor (AF) 647 or FITC. After incubation at 4°C for 30 min, cells were pelleted and resuspended in 2.5-5 mL AT1R staining buffer, followed by the addition of 250-500 µL anti-647 or anti-FITC microbeads (Miltenyi). Yeast cells were incubated with microbeads for 20 min at 4°C and then pelleted, resuspended in staining buffer, and added to 2 or 3 LS columns (Miltenyi). The columns were then washed with staining buffer and bound yeast cells were eluted in 5 mL staining buffer. From these elutions, ∼1 x 10^8^ cells were pelleted for FACS and stained with FLAG-tagged angiotensin II type 1 receptor (*5*) for 45 min at 4°C. Following this, yeast cells were again pelleted and stained with fluorescently labeled anti-FLAG antibody and fluorescently labeled anti-HA antibody for 20 min at 4°C, alternating between FITC, 647, and 488 labeled antibody for each AHEAD cycle to avoid selection for dye binding. FACS was performed with a Sony SH800 using a 100 µm Sony Sorting Chip. Over the course of nanobody AT110 affinity maturation, cells were grown for a total of ∼400 divisions. During that time, eight cycles of AHEAD were performed.

In preparation of next-generation sequencing, p1 plasmid was extracted, as previously described (*6*), from yeast cultures after the FACS step of each AHEAD cycle. PCRs were performed with Q5 Master Mix (New England Biolabs) and primers NGS_p1_F and NGS_p1_R. Following PCR reactions, samples were PCR purified (Qiagen). Amplicon sequencing was performed by the MGH CCIB DNA Core and the resulting sequences were analyzed with Geneious Prime.

### Radioligand binding assay for anti-AT1R nanobodies

Nanobodies were purified from the periplasm of *Escherichia coli* by Ni-NTA affinity chromatography and dialyzed into buffer consisting of 20 mM HEPES pH 7.4, 100 mM NaCl. Since AT110 and its derivatives allosterically increase the affinity of agonists for the AT1R, the effect of serially diluted nanobodies on the binding of the inverse agonist [^3^H]-olmesartan was assessed in the absence versus the presence of an ∼IC_20_ concentration of the low-affinity agonist, TRV055. Purified wild-type AT1R (75 ng) (*6*) was added to 2.5 nM [^3^H]-olmesartan (American Radiolabeled Chemicals, Inc), the indicated concentration of nanobody, and either assay buffer (20 mM HEPES pH 7.4, 100 mM NaCl, 0.01% lauryl maltose neopentyl glycol, and 0.1% bovine serum albumin) or 1 μM TRV055. The final reaction volumes were 200 μL, with single replicates for each condition. Assay buffer consisted of 20 mM HEPES pH 7.4, 100 mM NaCl, 0.01% lauryl maltose neopentyl glycol, and 0.1% bovine serum albumin. After a 90 minute incubation at room temperature, reactions were harvested onto GF/B glass-fiber filter paper using a 96-well harvester (Brandel) and quickly washed three times with cold 20 mM HEPES pH 7.4, 100 mM NaCl. The fraction of [^3^H]-olmesartan bound in the presence versus the absence of TRV055 was determined at each nanobody concentration. Data from three independent experiments were fit to a one-site model in GraphPad Prism.

### Engineering a stronger secretory leader sequence for the second-generation AHEAD 2.0 system

One of the parts responsible for higher nanobody display levels in AHEAD 2.0 is a mutated app8 secretory leader, which we selected from an error-prone PCR library of app8 (**Fig. S2**). Nanobody AT110 was fused to an app8 secretory leader (*32*) and cloned into a nuclear CEN/ARS plasmid containing either a pREV1, pSAC6, pRPL18B or a pTDDH3 promoter (*32*). Nanobody display from the pSAC6 promoter was determined to be most similar to expression from p1 with the pGA promoter. This plasmid, dubbed pAW258 was then used as template for preparing a library of mutated app8 secretory leader sequences. app8 was amplified by error prone PCR (GeneMorph II, Agilent) using primers AW_Sac6_mut_F and AW_Sac6_mut_R and cloned back into pAW258 using Gibson assembly. The library size was determined to be 2×10^7^ by counting colonies on serially diluted antibiotic selection plates. Twelve single clones were picked and sequenced. All tested clones had intact reading frames while the average number of mutations per clone was The library was then transformed into EBY100 cells, induced through growth in 2% galactose as the sole sugar source for 1-2 days, and subjected to three rounds of FACS selection for strong HA tag display, as the HA tag is fused to AT110 and acts as a surrogate for AT110 display level. In each round, the top 0.05% expressing cells as determined by anti-HA signal were sorted (**Fig. S2C**). After the third round, the cell population was plated, 48 single colonies were picked and screened for nanobody display. The 12 clones that displayed the highest levels of nanobody expression were sequenced. Three different mutations were discovered, namely, V10A, F48V and D28G. Those mutations were reintroduced to plasmid p258 and assayed for their effect on AT110 display level. While mutations F48V and D28G did not confer any increase in display levels, mutation V10A increased nanobody display by ∼90% (**Fig. S2C**). App8 with V10A was dubbed app8i1 and used in the second-generation AHEAD 2.0, along with other modifications described above.

### Affinity maturation of Nb.b201 and Lag42 using the second-generation AHEAD 2.0 system

HSA (Sigma) was directly labeled with AF647. 6XHis tagged GFP was expressed in *E. coli* and purified using Ni-NTA agarose (ThermoFisher). Nb.b201 was amplified using PCR from pYDS-Nb.b201 (*19*) and cloned into pAW240. Lag42 was synthesized as a gBlock (IDT DNA technologies) and cloned into pAW240. The resulting plasmids were linearized using ScaI, transformed into yAW301, and plated as described above (see **Materials and Methods** section “cloning nanobodies into AHEAD”). A single colony was picked into SC-HLUW, grown to saturation, pelleted, induced in SC-HLUW with 2% galactose replacing glucose as the sole sugar source for ∼24 hours, and labeled with 50 nM HSA-AF647 (for Nb.b201 cells) or 200 nM GFP-AF647 (for Lag42 cells) and 1 µM mouse anti-HA antibody for 1 hour at 4°C in binding buffer (20 mM HEPES pH 7.5, 150 mM NaCl, 0.1% BSA, 0.2% maltose). Cells were washed with binding buffer and incubated with 0.5 µM polyclonal goat anti-mouse AF488-labeled antibody (ThermoFisher) for 15 minutes. Cells were washed again with binding buffer and sorted (SONY SH800) for increased affinity for HSA or GFP (**Fig. S3**) into a tube containing 2 ml SC-HLUW. Cells were incubated with shaking for 1-2 days at 30°C and subjected to the next AHEAD cycle.

Over 4 cycles of AHEAD (for Nb.b201) or 6 cycles of AHEAD (for Lag42), the selection stringency was increased by reducing the concentration of HSA or GFP as indicated (**Fig. S3**). During each FACS step, ∼2 x 10^7^ cells were used for sorting out 200-1000 cells. After AHEAD, nanobodies were amplified from p1 and either sequenced directly or subcloned into a plasmid to isolate individual clones for sequencing and further characterization.

### Isolation of anti-RBD nanobody parents

In order to isolate RBD-binding nanobodies, two initial rounds of magnetic-activated cell sorting (MACS) were performed using a synthetic yeast-displayed library of nanobodies (*19*) followed by two rounds of FACS. *S. cerevisiae* containing the library were grown in tryptophan dropout media (US Biological) with 2% glucose for 1 day and then, expression and display was induced in tryptophan dropout with 2% galactose for 2 days. For the first round, 1 x 10^10^ yeast were centrifuged and resuspended in a ‘pre-clearing’ solution of 4.5 mL of binding buffer, 500 µl of anti-PE microbeads (Miltenyi), and 200 nM streptavidin-PE (BioLegend). After incubation for 40 min at 4°C, yeast cells were passed through an LD column (Miltenyi) to remove cells interacting with microbeads or streptavidin. Yeast cells that flowed through the column were collected, centrifuged, resuspended in a ‘staining solution’ consisting of 2 mL binding buffer with 1 µM SARS-CoV-2 RBD and 250 nM streptavidin-PE and incubated for one hour at 4°C. After incubation, yeast cells were centrifuged, resuspended in a ‘secondary solution’ of 4.5 mL binding buffer and 500 µL anti-PE microbeads, and incubated an additional 15 min at 4°C. These yeast cells were then centrifuged, washed with binding buffer, and passed into an LS column (Miltenyi). The LS column was washed with 7 mL of binding buffer and remaining yeast were eluted in 5 mL of binding buffer, centrifuged, and resuspended in 5 mL tryptophan dropout media for expansion. The second round of MACS was performed similarly to the first but starting with 1 x 10^9^ yeast and substituting PE-labeled streptavidin with FITC-labeled streptavidin and anti-PE microbeads with anti-FITC microbeads. Additionally, volumes of the pre-clearing and secondary solutions were reduced 5-fold and the staining solution by 2-fold.

For the first round of FACS, 1 x 10^8^ induced cells were stained with 1 µM of directly AF488-labeled SARS-CoV-2 RBD and 0.5 µM anti-HA AF647 (Cell Signaling Technology) antibody, to visualize expression, for 1 hr at 4°C. These cells were then centrifuged, resuspended in 2 mL binding buffer, and sorted. In total, 35,000 cells from 11,431,000 were collected and expanded. The second round of FACS was performed with similar conditions to the first; however, the RBD was labeled with AF647, anti-HA with AF488, and the concentration of RBD was reduced to 150 nM. For the second round, 104,000 cells were collected from 2,330,000 sorted. These cells were expanded in culture and then plated on tryptophan dropout media to isolate single clones. Twenty-four colonies were picked, cultured, and induced. Each culture was screened for binding by staining ∼1 x 10^6^ cells with 200 nM of 647- and 488-labeled RBD along with 488- and 647-labeled anti-HA antibody, respectively, and binding reactions were evaluated using a BD Accuri flow cytometer. Promising clones were selected as parents for AHEAD experiments.

### FACS selection for improved RBD binders using the improved second-generation AHEAD system

After cloning the RBD-binding parent nanobodies into AHEAD (see **Materials and Methods** section “cloning nanobodies into AHEAD”), initial cultures (50 mL SC-HLUW) were grown to saturation and optionally passaged once or twice into 50 mL SC-HLUW at a ratio of 1:1000 to prolong diversification of the nanobodies before the first AHEAD cycle. Upon final saturation (1-2 days), cells were pelleted and resuspended in SC-HLUW with 2% galactose replacing glucose as the sole sugar source for induction. Induction was done for 24 hours at room temperature with shaking at 250 rpm. Cells were collected, washed in binding buffer (20 mM HEPES pH 7.5, 150 mM NaCl, 0.1% BSA, 0.2% maltose) and incubated with 1 µM mouse anti-HA antibody labeled and SARS-CoV-2 RBD directly labeled with AF647. Cells were then washed and incubated with 0.5 µM goat anti-mouse AF488-labeled antibody (ThermoFisher) for 15 minutes. The cells were subjected to FACS (SONY SH800) whereby 200-500 cells were collected into a culture tube containing 3 mL SC-HLUW out of ∼2 x 10^7^ sorted cells. Each subsequent cycle of AHEAD involved growing the 3 mL of sorted cells with shaking at 30°C until saturation (1-2 days), induction with 2% galactose as the sole sugar source for ∼24 hours at room temperature with shaking at 250 rpm, washing of cells in binding buffer, incubation with anti-HA and labeled RBD, washing steps to remove unbound RBD, and FACS sorting into 3 mL SC-HLUW. RBD concentrations were diminished from cycle to cycle while the stringency of washes increased. After several rounds of AHEAD, nanobodies were amplified from p1 and either sequenced directly or subcloned into a plasmid to isolate individual clones for sequencing and further characterization.

### Nanobody-Fc fusion purification

Nanobodies targeting RBD were expressed and secreted as Fc fusions by cloning into pFUSE-hlgG1-Fc2 (Invivogen) using the NcoI and EcoRI restriction sites or by Gibson assembly. For each nanobody-Fc fusion, 100 mL of Expi293 cells were transfected with 90-150 µg of plasmid. After 1 day, cells were enhanced with 3 mM valproic acid and 0.45% glucose. Cell supernatants were harvested 4 days after transfection. Before purification, supernatants were treated with benzonase nuclease and protease inhibitor, then passed through a 0.22 µm filter. Nanobody-Fc fusion supernatants were passed over a column with 4 mL protein G resin (ThermoFisher), which was then washed with 40 mL of HBS, eluted with 100 mM citrate (pH 3) and then neutralized to pH 7 with concentrated HEPES (pH 8). Nanobody-Fc fusions were then dialyzed twice with HBS (pH 7.5).

### On-yeast EC_50_ measurements

To determine nanobody affinities for their targets as surface-displayed proteins, individual nanobody sequences were cloned into plasmid p253, a plasmid for galactose-inducible expression of nanobodies for surface display in EBY100 cells. Plasmids were transformed into EBY100, induced in appropriate dropout media with 2% galactose as the sole sugar source for ∼24 hours at room temperature, washed with binding buffer (20 mM HEPES pH 7.5, 150 mM NaCl, 0.1% BSA, 0.2% maltose), and labeled with biotinylated antigen across a range of concentrations as well as with 1 μM mouse anti-HA antibody for 1 hour at 4°C. Cells were then washed and incubated 0.5 μM with goat anti-mouse AF488-labeled antibody (ThermoFisher) and streptavidin conjugated PE (Abcam) for 15 minutes. After additional washing, fluorescence was measured using an Attune flow cytometer (Life Technologies). Antigen binding (PE signal) was recorded only for cells that express the nanobody, namely cells in populations showing anti-HA staining signal. Average PE signal at each antigen concentration was determined and used to fit a one-site model in GraphPad Prism in order to determine the EC_50_. In all cases, volumes and number of cells used were chosen to accommodate doing assays in 96-well format and to avoid ligand depletion. Binding was measured in triplicate for each antigen concentration.

### Surface plasmon resonance

SPR was performed using a Biacore T200 (Cytiva). Nanobody Fc fusion proteins were immobilized to a protein A sensor chip (Cytiva) at a capture level of approximately 85–255 response units (RU). Binding experiments with dilutions of RBD were performed in running buffer (10 mM HEPES pH 7.5, 150 mM NaCl, 0.05% Tween). RBD dilutions were injected at a flow rate of 30 µL/min with a contact time of either 160 seconds or 280 seconds and a dissociation time of 600 seconds or 900 seconds. After each cycle, the protein A sensor chip was regenerated with 10 mM glycine-HCl Ph Kinetic data were double reference subtracted and fit to a 1:1 binding model. For samples in which the on or off rates could not be determined, data were fit to a steady-state affinity model.

### SARS-CoV-2 and VSV-G Lentivirus Production

To generate lentivirus pseudotypes, the SARS-CoV-2 virus spike protein with the last 27 amino acids deleted (Genbank ID: QJR84873.1 residues 1-1246) was cloned into a pCAGGS vector and modified to include at its C-terminal tail the eight most membrane adjacent residues of the cytoplasmic domain of the HIV-1 envelope glycoprotein (NRVRQGYS). Pseudotypes were packaged by transfecting HEK293T cells (ATCC CRL-11268) using lipofectamine 3000 (Invitrogen) with SARS-CoV-2 S in pCAGGS or VSV G in pCAGGS (as previously described (*34*)), in addition to a packaging vector containing HIV Gag, Pol, Rev, and Tat (psPAX2, provided by Didier Trono, Addgene # 12260), and a pLenti transfer vector containing GFP (pLenti-EF1a-Backbone, a gift from Feng Zhang (*35*), Addgene plasmid #27963). After 18 hours, the transfection medium was removed from cells and replaced with DMEM containing 2% (v/v) fetal bovine serum (FBS) and 50 mM HEPES. Cells were incubated at 34°C, before the supernatant was then harvested at 48 and 72 hours, centrifuged at 3000 x g, and filtered through a 0.45 µm filter. The filtered supernatant was then concentrated by layering on a 10 % (v/v) sucrose cushion in Tris Buffered Saline (50 mM Tris-HCl pH 7.5, 100 mM NaCl, 0.5 mM EDTA) and spun at 10,000 x g for 4 hours at 4°C. The viral pellet was resuspended in Opti-MEM containing 5% (v/v) FBS and stored at −80°C.

### Pseudotype Neutralization Experiments

Nanobodies or PBS alone were pre-incubated with SARS-CoV-2 or VSV G lentivirus for 60 minutes at 37°C in a mixture that also contained 0.5 µg/ml polybrene. Mixtures were then added to HEK293T cells overexpressing human ACE2 (a gift from Dr. Huihui Mou and Dr. Michael Farzan, Scripps Research). After 24 hours the virus medium was removed and replaced with DMEM containing 10% (v/v) FBS, 5% (v/v) Pen/Strep, and 1 µg/mL puromycin. The percentage of GFP positive cells was determined by flow cytometry with an iQue Screener PLUS (Intellicyt) 48 hours after initial infection. Percent relative entry was calculated using the following equation: Relative Entry (%) = (% GFP positive cells in nanobody well)/(%GFP positive cells in PBS alone well). Percent neutralization was calculated using the following equation: Neutralization (%) = 1-(% GFP positive cells in nanobody well)/(% GFP positive cells in PBS alone well). All experiments were performed twice in triplicate.

### ACE2 competition assays using biolayer Interferometry

Biolayer interferometry (BLI) experiments for ACE2 competition assays were performed with an Octet RED96e (Sartorius). For ACE2 competition assays, biotinylated SARS-CoV-2 RBD was loaded onto streptavidin (SA) sensors (ForteBio) at 1.5 µg/ml for 80 seconds. After a baseline measurement was obtained, nanobodies were associated at 250 nM for 300 seconds, followed by an association with ACE2-Fc at 250 nM for 300 seconds. Representative results of two replicates for each experiment are shown.

### Recombinant RBD expression and purification

Two preparations of the receptor-binding domain (RBD) of SARS-CoV-2 S (GenBank ID: QHD43416.1, residues 319-541) were used in this study. For ACE2 competition assays, RBD was cloned into the pHLsec vector (Ref. PMID: 17001101). The construct contains an N-terminal 6xHis tag, a TEV protease site, and a BirA ligase site followed by a 7-residue linker. RBD was produced by transfecting HEK293T cells grown in suspension and harvested after 5 days and was purified by reverse nickel affinity purification. The RBD was then biotinylated with BirA ligase and again purified using reverse nickel affinity purification to remove the BirA ligase, followed by size exclusion purification on a Superdex 200 Increase column (GE Healthcare). For all other experiments including AHEAD evolution of anti-RBD nanobodies and their characterization, a pVRC8400 plasmid containing RBD was used for RBD expression (*36*). The construct (pVRC8400-RBD) has RBD fused to a C-terminal HRV C3 protease cleavage site followed by an 8XHis tag and a streptavidin binding peptide (SBP). 0.5 mg of plasmid was transfected into HEK293T cells and grown in suspension for 3 days. Culture media was then dialyzed against PBS overnight and RBD was purified using Ni-NTA agarose (ThermoFisher). The eluted His-tagged RBD was then incubated with biotin-tagged HRV C3 protease (Sigma) and passed through a streptavidin-agarose column to deplete the protease and the 8XHis-SBP peptide. The eluate was collected and analyzed on a denaturing SDS-PAGE gel to confirm purity of the RBD.

### Recombinant ACE2-Fc fusion protein expression and purification

To generate a recombinant ACE2 Fc-fusion protein, we cloned the ectodomain of human ACE2 (GenBank ID: BAB40370.1 residues 18-740) with a C-terminal Fc tag into the pVRC8400 vector containing the human IgG1 Fc. We transfected the construct (pVRC8400-hACE2) into Expi293F™ cells using an ExpiFectamine™ transfection kit according to the manufacturer’s protocol. The supernatant was harvested after 5 days and purified using a MabSelect SuRE Resin (GE Healthcare) followed by size exclusion purification on a Superose 6 Increase column (GE Healthcare). The supernatant was harvested 5 days after transfection and purified with a CaptrueSelect KappaXL Affinity Matrix (Thermo Scientific) followed by size exclusion chromatography on a Superdex200 Increase column (GE Healthcare).

### Cloning a 200,000-member nanobody library into AHEAD2.0

A computationally designed 200,000-member naïve nanobody CDR3 library was synthesized as an oligonucleotide pool of CDR3 sequences by Agilent, as previously reported (*29*). To clone this CDR3 library into AHEAD, we first made plasmid pAW240-NbCM, which encodes the nanobody scaffold with fixed CDR1 and CDR2 sequences but with CDR3 replaced with a NotI restriction site. The CDR3 oligo library was then inserted into pAW240-NbCM using the NotI site and ligation. Transformation of the ligated plasmid products into *E. coli* resulted in ∼10^8^ transformants. 200 µg of plasmid DNA was then prepared, linearized with ScaI (NEB) and transformed into strain yAW301 by scaling up the process described in **Materials and Methods** section “cloning nanobodies into AHEAD” 100-fold. The total transformation resulted in ∼10^7^ transformants such that the 200,000-member library was covered 50X.

### Selection and affinity maturation of a GFP binding nanobody from a computationally designed naïve nanobody library encoded on AHEAD

A 10 mL saturated culture of yAW301 cells expressing the computationally deigned 200,000-member naïve nanobody library was induced in SC-HLUW with 2% galactose replacing glucose as the sole sugar source. Induction was done for 24 hours at room temperature with shaking at 250 rpm. Cells were collected, washed in binding buffer (20 mM HEPES pH 7.5, 150 mM NaCl, 0.1% BSA, 0.2% maltose), and first subjected to negative selection against streptavidin binders by MACS. Specifically, cells were incubated for 1 hour at 4 °C with 0.5 µM streptavidin-conjugated FITC, washed, and incubated with anti-FITC microbeads (Miltenyi). Cells were washed again and passed through an LD column to deplete streptavidin binders. Recovered cells eluted from the column were incubated with 200 nM GFP-biotin and 1 µM mouse anti-HA antibody. Cells were washed and incubated for 15 minutes in binding buffer containing 0.5 μM goat anti-mouse AF488-labeled antibody (ThermoFisher) and streptavidin-conjugated AF647. Cells were washed again and subjected to the first cycle of FACS for GFP binders. In the following cycles of AHEAD, cells were incubated with GFP directly labeled with AF647 (**Fig. S8**). These cycles of AHEAD followed the same process for nanobody evolution using AHEAD 2.0 described above.

## Acknowledgments

We thank W. Capel for assistance with nanobody purifications. We thank Z. Zhong, C. Carlson, T. Loveless, A. Banks, and other members of the Liu and Kruse groups for experimental assistance, materials, and thoughtful discussions. We thank G. Arzumanyan for the pGA promoter discovered in his OrthoRep continuous protein evolution experiments unrelated to this study. This work was funded by NIH 1DP2GM119163, NIH NIGMS 1R35GM136297, the Moore Inventor Fellowship, and the UCI COVID-19 Basic, Translational and Clinical Research Fund to C.C.L.; NIH DP5OD021345 and a Vallee Scholars Award to A.C.K.; NIH NIAID R01AI146779 and a Massachusetts Consortium on Pathogenesis Readiness (MassCPR) grant to A.G.S.; training grants NIGMS T32GM007753 for B.M.H. and T.M.C. and T32AI007245 for J.F.; and NIH NCI 1R01CA260415 to C.C.L., A.C.K., and D.S.M.

## Author contributions

All authors contributed to experimental design and data analysis. A.W., C.M., A.C.K., and C.C.L. were responsible for the conception of AHEAD. A.W. and K.M.N. carried out experiments establishing the first-generation of AHEAD and made improvements to reach the second-generation AHEAD system. C.M. carried out AHEAD experiments for the evolution of anti-AT1R nanobodies and selected parent anti-SARS-CoV-2 for evolution using AHEAD. A.W., J.C., and M.H.H. carried out AHEAD experiments for the evolution of anti-GFP, anti-HSA, and anti-SARS-CoV-2 nanobodies. A.W., C.M., M.S.A.G., S.C., and L.M.W. characterized the activities of evolved nanobodies in binding assays (A.W., C.M., and L.M.W.), SPR measurements (C.M. and M.S.A.G.), neutralization assays (S.C.), and ACE2 competition assays (S.C.). J.F., B.M.H., T.M.C., and A.W. were responsible for the expression of RBD used throughout this study. J.-E.S. and D.S.M. were responsible for computational design aspects for the naïve ∼200,000-member nanobody library and A.W. inserted that library into AHEAD. A.C.K. and C.C.L. oversaw all aspects of the project, D.S.M. supervised computational nanobody library design, J.A. supervised neutralization and ACE2 competition assays, and A.G.S. supervised the preparation of RBD. A.W., C.M., A.C.K., and C.C.L. wrote the manuscript with input and contributions from all authors.

## Data and materials availability

Correspondence and requests for materials should be addressed to ACK and CCL.

## Supplementary Information for

### Supplementary Text

#### Functional display of AT110 and 4-4-20 from AHEAD1

When we first examined whether functional surface display was feasible from p1, by testing the 4-4-20 scFv and the AT110 nanobody, we used the wild type DNAP for replicating p1 (**Fig. 1C**). However, when we transitioned to using the error prone polymerase, TP-DNAP1-4-2 the functional display levels dropped significantly (**Fig. S2D**) resulting from a decline in p1’s copy number as described previously (*3, 4*). The weak binding signal limited FACS selections, thus we overcame these limitations by MACS, and later on by designing AHEAD2.0 as described below.

#### Enhancing nanobody expression by promoter and secretory leader engineering

Our evolution of improved AT110 nanobodies was performed using AHEAD 1.0, a system that conferred low nanobody expression. As discussed in the main text, in that experiment, insufficient nanobody expression was overcome by implementing a MACS enrichment protocol for cells that had the highest levels of nanobody expression. However, this introduces additional days to the AHEAD cycle because of the time required for MACS and because cells from the (N-1)^th^ cycle of AHEAD need to populate larger culture volumes in the N^th^ cycle of AHEAD in order to have sufficient numbers of cells for FACS after a selective MACS step for high expressers. Therefore, the requirement for MACS enrichment, while seemingly simple, significantly eroded the goals of AHEAD by both extending the time for an AHEAD cycle and reducing parallelizability, given the increased difficulty of working with large cultures. To solve these issues, we sought to increase expression of nanobodies to the point where MACS pre-enrichment for expression was not required. To obtain higher expression of yeast displayed nanobodies from p1, we targeted the promoter and the secretory leader sequence for engineering. Fortunately, in an unrelated OrthoRep continuous experiment performed (G. Arzumanyan, unpublished), three expression-enhancing promoter mutations were discovered in the p1-specific p10B2 promoter, which was the strongest promoter for driving expression from p1 at that time (*4*). We tested all possible combinations of the three promoter mutations, which revealed a pair of mutations, G5A and G34A (upstream to the start codon) that together increased nanobody expression by 3.5-fold. The new promoter containing those mutations was dubbed pGA. To improve display efficiency further, we chose to engineer the strongest known alpha factor secretory leader, app8 (*8*). As described in **Materials and Methods**, an error-prone PCR library of app8 variants was screened for mutants that raised display levels, leading to app8i1. Finally, a poly adenosine tail (A75) followed by a self-cleaving hammerhead ribozyme (RZ) was added to the nanobody expression and display cassette on p1 in order to confer an additional increase in expression following our past work (*4*). With the improved system, productive FACS selections for antigen binding by displayed nanobodies could be implemented after every cycle of cell propagation in small culture volumes of 2-5 mL. This led to an AHEAD 2.0 cycle time of only 3 days with high parallelizability.

Interestingly, many of the AHEAD 2.0 evolution experiments described in this paper led to fixation of additional mutations in the pGA promoter as well as the app8 leader. Although these mutations have not been characterized in this study, they likely fixed because they increased nanobody display efficiency and robustness and will be useful for future versions of AHEAD.

#### Specific considerations for gating during FACS in AHEAD experiments

In yeast display antibody evolution, one typically gates for target-binding normalized to the nanobody display level. To measure nanobody display levels, we used a human influenza hemagglutinin (HA) tag fused to the nanobody. We then gated on the ratio of [target (*i*.*e*. RBD) binding]:[HA tag binding by a fluorescently labeled anti-HA antibody]. In other words, during selection for better target binding, we gate along a slope on the FACS plot where the Y-axis is HA level and X-axis is target-binding level (see **Fig. S5** for examples). However, since AHEAD hypermutates the entire content of p1, it is possible to obtain mutations in the HA tag that disable it, creating cells that have seemingly strong target binding per nanobody displayed if sorted exclusively for cells with a high [target binding]:[HA signal] ratio. This can lead to the gradual fixation of cheaters that actually bind the target weakly but were selected because they also have disabled HA tags. To overcome this issue, it is important to gate with a strict floor on HA signal rather than solely on the [target binding]:[HA signal] ratio. In our experiments, we found that if the floor for HA signal was set to include only the top ∼20% of cells on the HA signal axis (Y-axis), we could maintain selection for target-binding throughout rounds without carrying over cheaters that mutated the HA tag.

Table S2. Information and activities for all anti-SARS-CoV-2 nanobodies characterized in this study. Please refer to file Table S2.xls.

---

**Fig S1.**
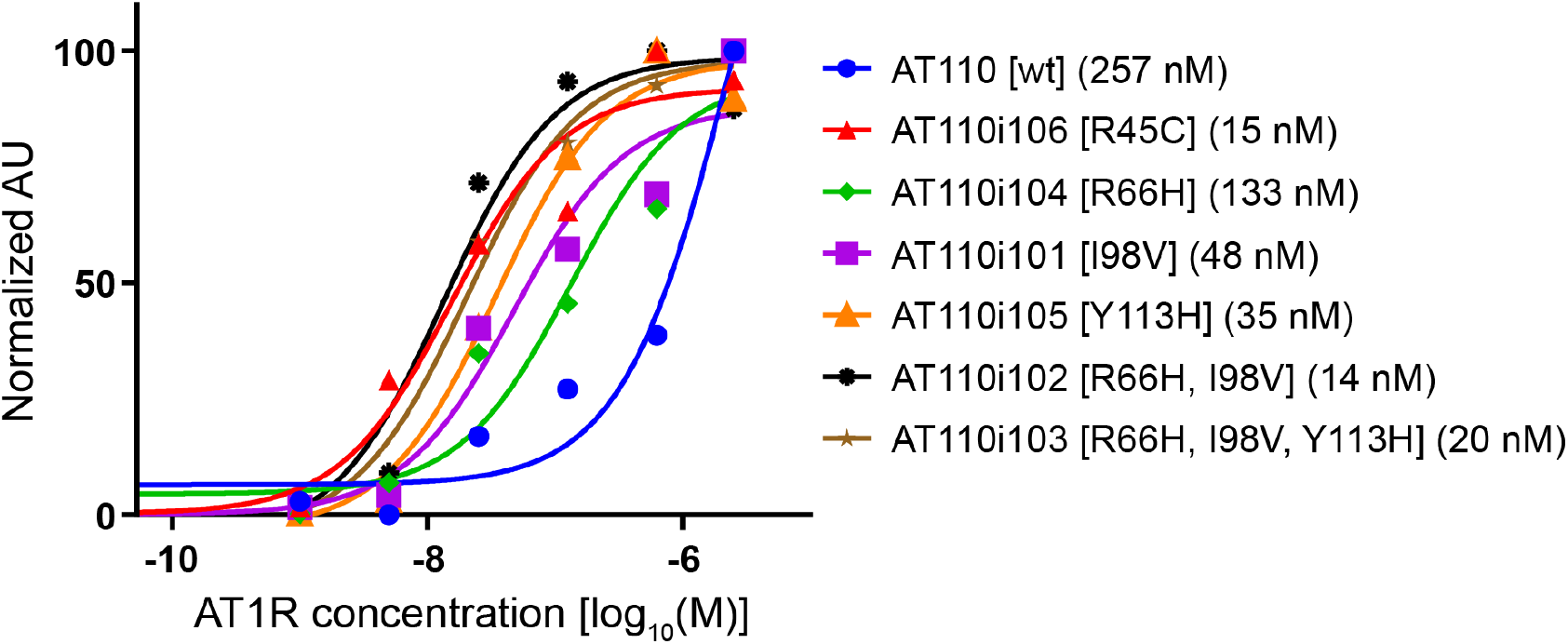
Contributions of individual mutations fixed during the evolution of AT110 by AHEAD. Affinity (EC_50_) of each nanobody for AT1R was determined by measuring binding at each concentration of AT1R (coupled to angiotensin II) in a single replicate and fitting the resulting binding curve. (See Materials and Methods.)

**Fig S2.**
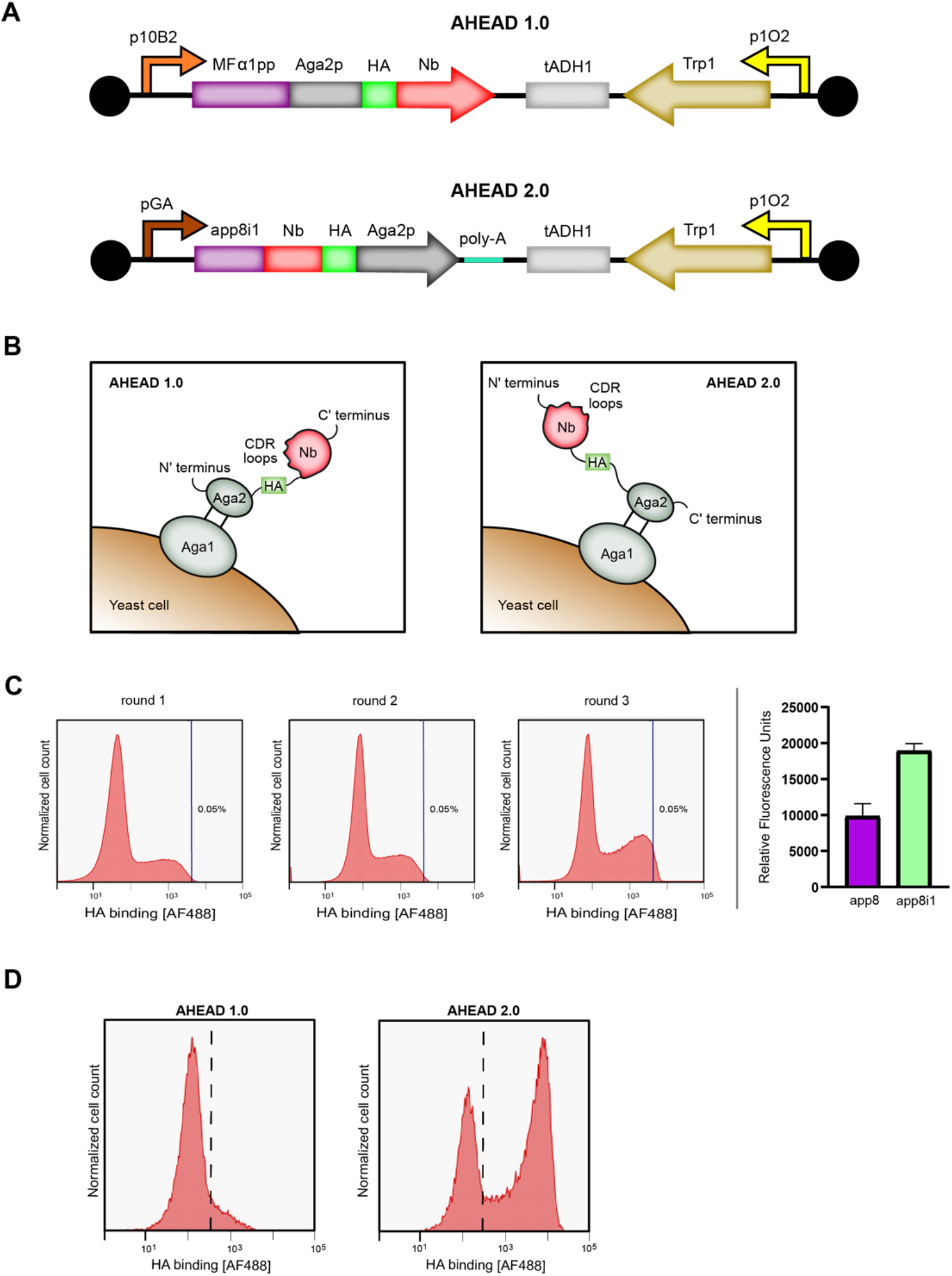
Optimization of antibody display in AHEAD. (**A**) Architectures of orthogonal p1 plasmids containing OrthoRep parts driving expression of nanobodies in the first-generation AHEAD 1.0 and improved second-generation AHEAD 2.0 systems. (**B**) Architectures for nanobody display in the first-generation AHEAD 1.0 and improved second-generation AHEAD 2.0 systems. (**C**) Selection of a new leader sequence for higher nanobody display. FACS plots showing the progressive enrichment of higher efficiency leader sequences across 3 rounds of selection (left panel). Nanobody display level using app8 compared to the selected app8i1 variant (right panel). N = 3, error bars represent ± s.d. (**D**) Increased functional nanobody expression using all AHEAD 2.0 parts as determined by FACS. The induced population in AHEAD 2.0 shows an ∼25-fold increase in nanobody display efficiency compared to AHEAD 2.0.

**Fig S3.**
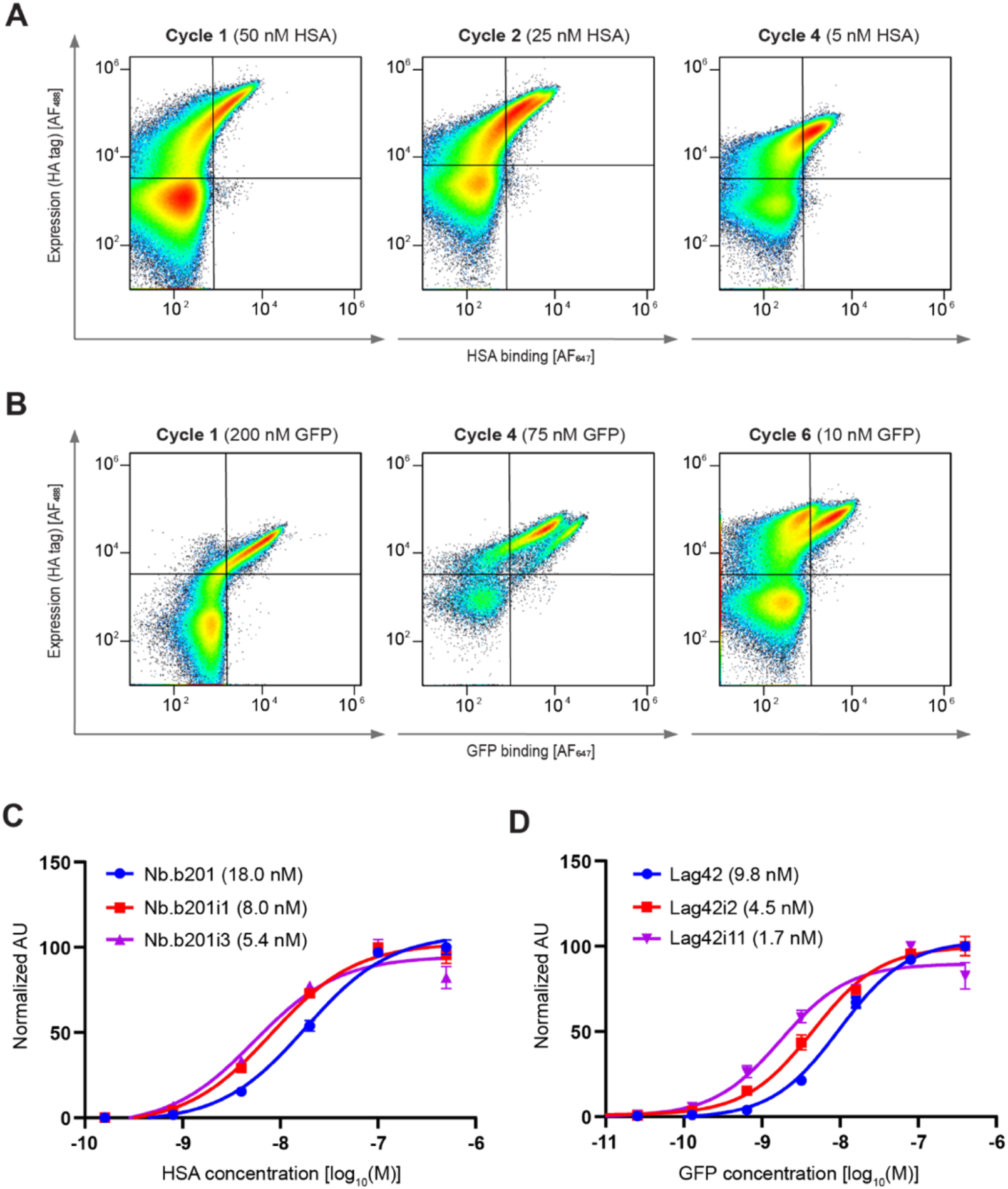
Evolution of anti-GFP and anti-HSA antibodies using an optimized second-generation AHEAD 2.0 system. (**A**) Selected FACS plots showing affinity maturation of Nb.b201 through AHEAD cycles. (**B**) Selected FACS plots showing affinity maturation of Lag42 through AHEAD cycles. (**C**) Affinities (EC_50_) of improved high-affinity anti-HSA nanobodies evolved using AHEAD. Binding by each concentration of HSA was determined in triplicate (error bars represent ± s.d.) and EC_50_s were determined by fitting each binding curve. (See Materials and Methods.) (**D**) Affinities (EC_50_) of improved high-affinity anti-GFP nanobodies evolved using AHEAD. Binding by each concentration of GFP was determined in triplicate (error bars represent ± s.d.) and EC_50_s were determined by fitting each binding curve. (See Materials and Methods.)

**Fig S4.**
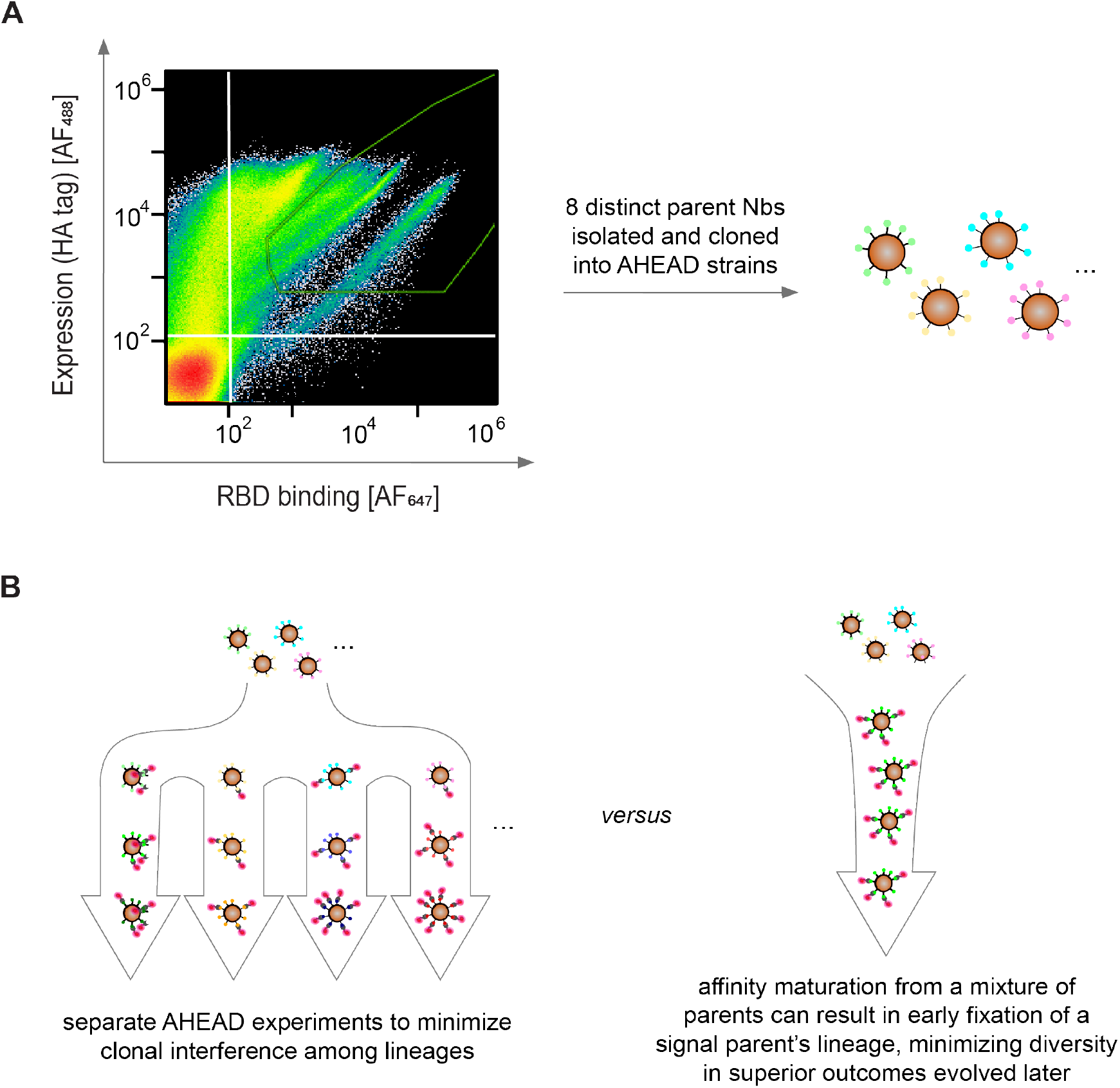
Isolation of parent anti-RBD nanobodies for AHEAD. (**A**) FACS plot showing enrichment of initial anti-RBD nanobody clones from a naïve nanobody library. The green polygon corresponds to the gate used for sorting. (**B**) Schematic showing the separation of parent clones into different AHEAD experiments to minimize clonal interference during affinity maturation.

**Fig S5.**
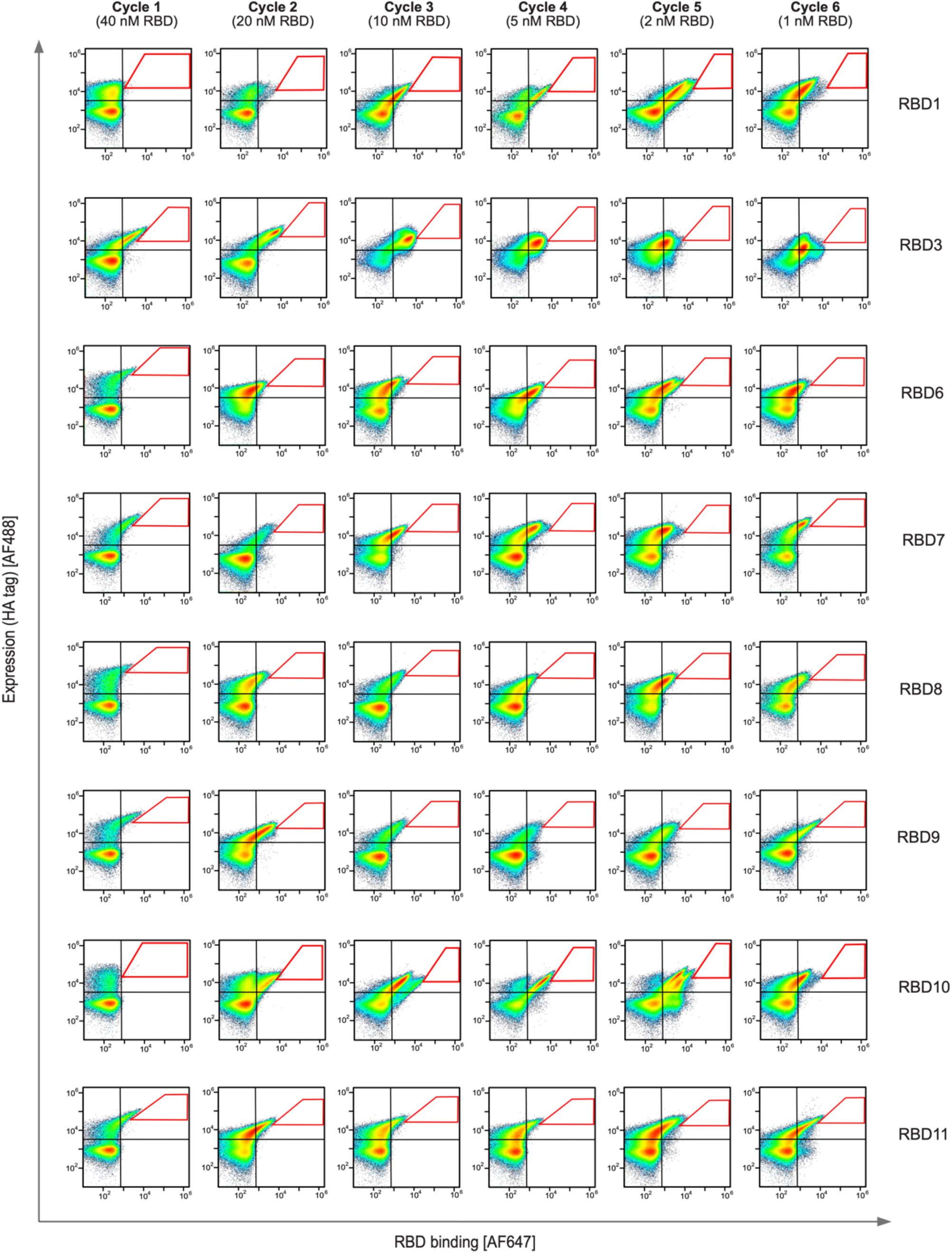
Selected FACS plots showing anti-RBD affinity maturation by cycles of AHEAD in 8 independent experiments starting from each of 8 parent clones identified from a naive nanobody library (see **Fig. S4**). Red polygons correspond to the gates used for sorting.

**Fig S6.**
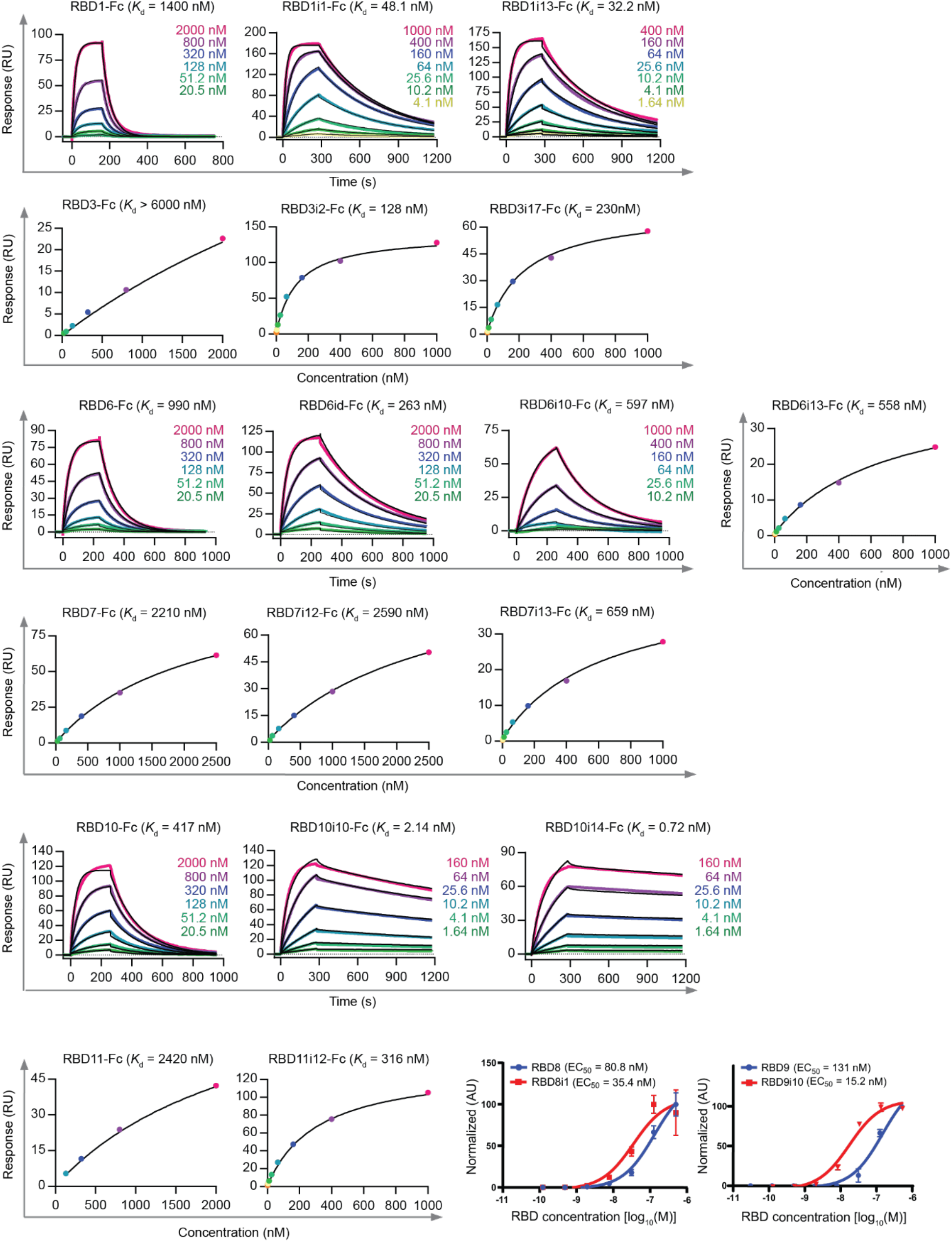
Affinities of anti-RBD nanobodies determined by surface plasmon resonance (SPR) or EC_50_ measurements. SPR or EC_50_ binding curves are shown for each anti-RBD nanobody characterized in this study. For EC_50_ affinities, binding by each concentration of RBD was determined in triplicate (error bars represent ± s.d.) and EC_50_s were determined by fitting each binding curve. (See Materials and Methods.). Steady state affinity fits were calculated for nanobodies for which the on and off rates could not be determined.

**Fig S7.**
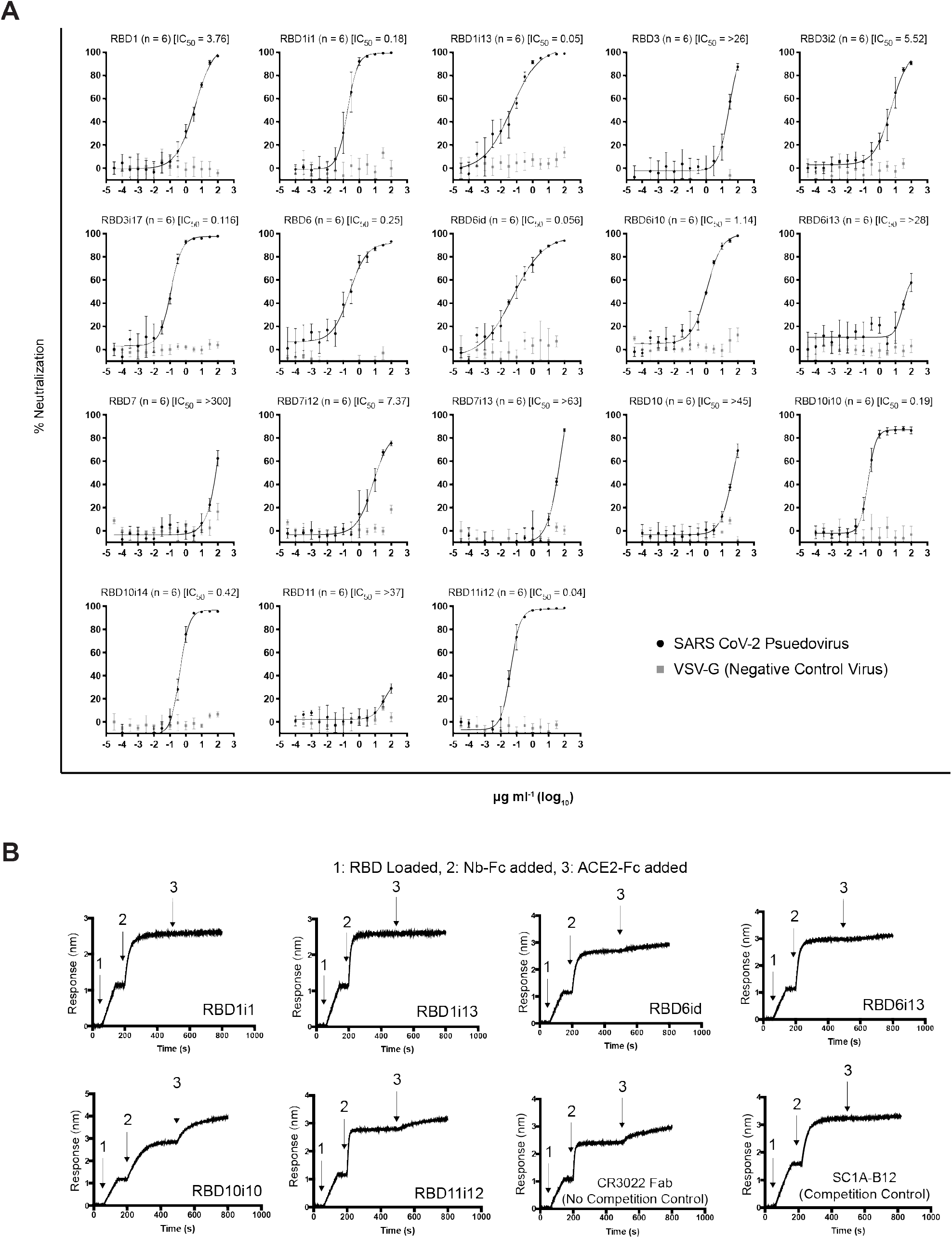
Neutralization assays and ACE2 competition assays for anti-RBD nanobodies evolved with AHEAD. (**A**) Neutralization plots for all anti-RBD nanobodies characterized in this study. Each nanobody concentration (X-axis) was tested in replicate. N = 6, error bars represent ± s.d. (**B**) Bio-layer interferometry (BLI) traces measuring ACE2 competition for anti-RBD nanobodies. CR3022 is an anti-RBD antibody that does not compete with ACE2 binding (no competition control) whereas SC1A-B12 is an anti-RBD antibody that competes strongly with RBD binding.

**Fig S8.**
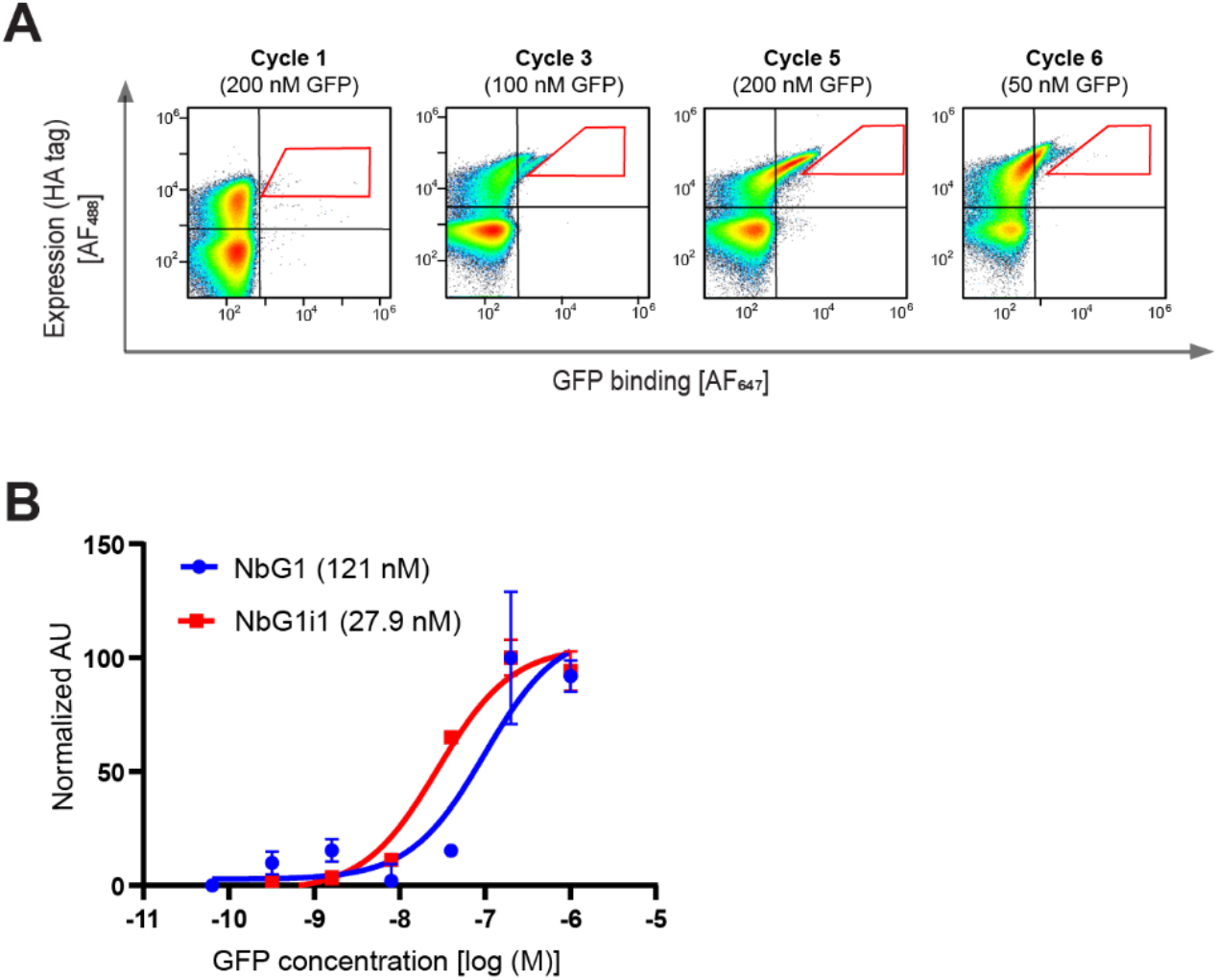
Evolution of an anti-GFP nanobody from a computationally-designed 200,000-member naïve nanobody library encoded on AHEAD. (**A**) Representative FACS plots showing enrichment of a GFP-binding clone from the nanobody library and subsequent emergence and fixation of a mutation that increases GFP binding across AHEAD cycles. (**B**) Affinity (EC_50_) of the AHEAD evolved anti-GFP nanobody, NbG1i1, isolated from AHEAD cycle 6 as compared to its parent, NbG1, that fixed in AHEAD cycle 3. Binding by each concentration of GFP was determined in triplicate (error bars represent ± s.d.) and EC_50_s were determined by fitting each binding curve. (See Materials and Methods.)

**Table S1.**
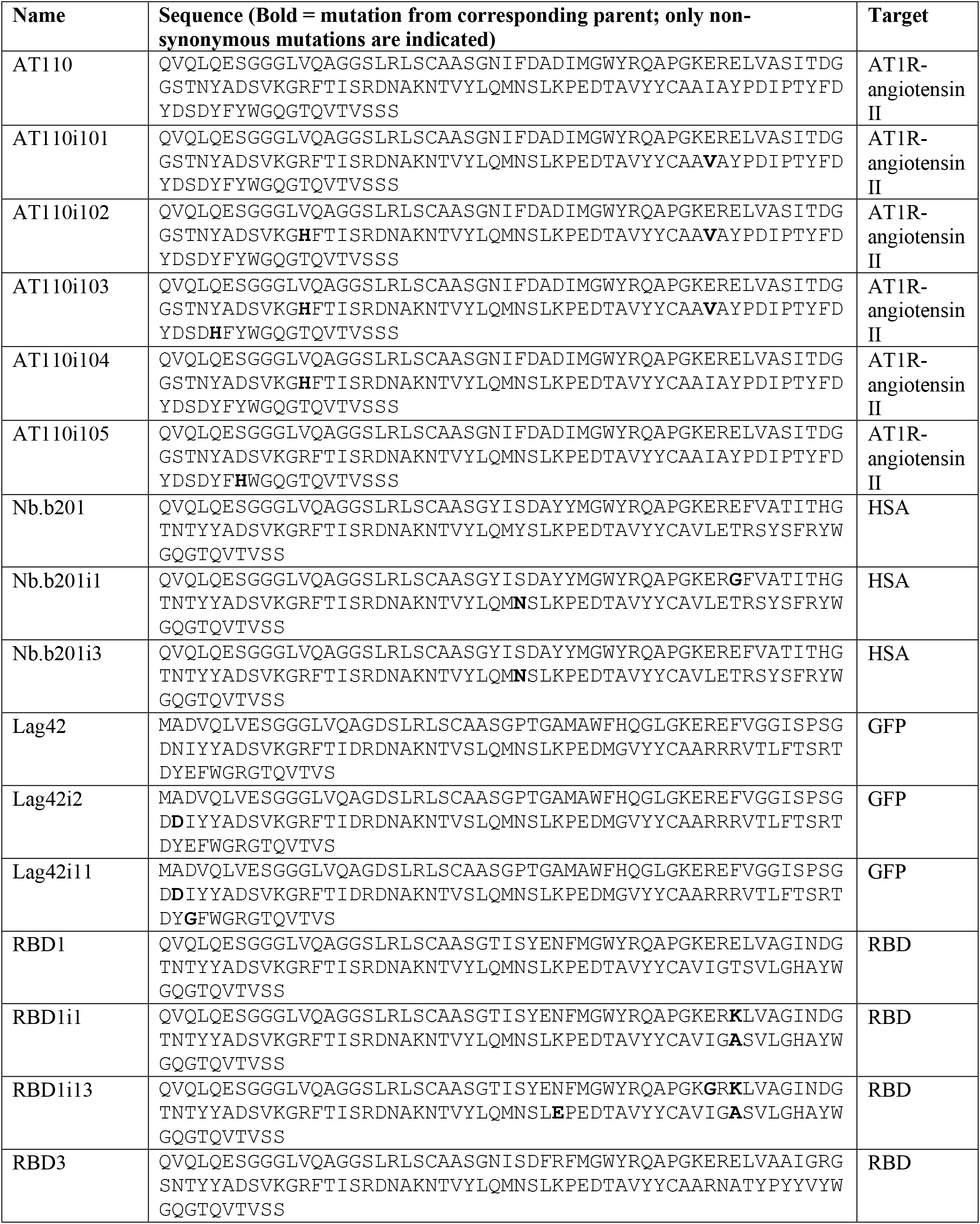
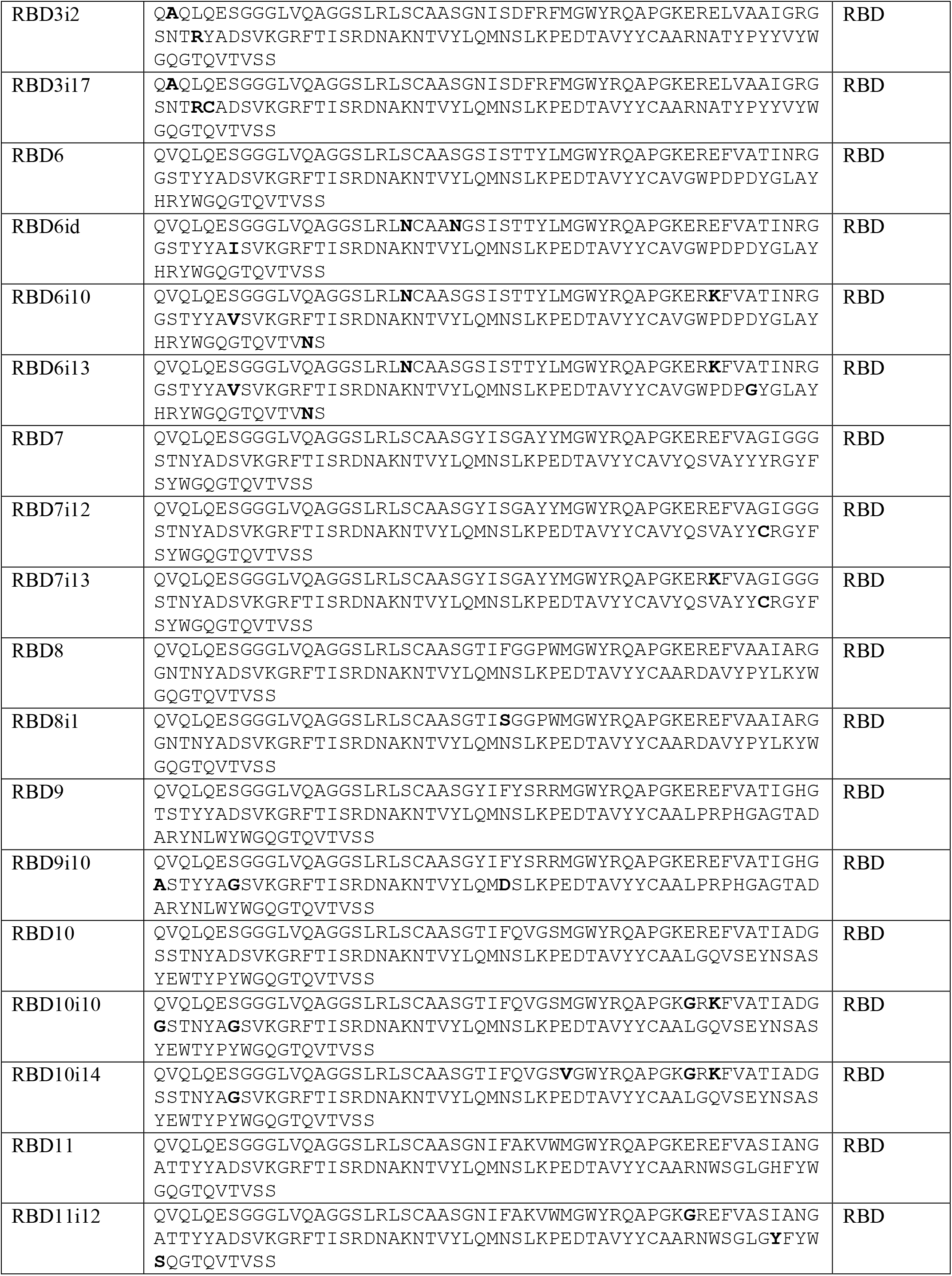
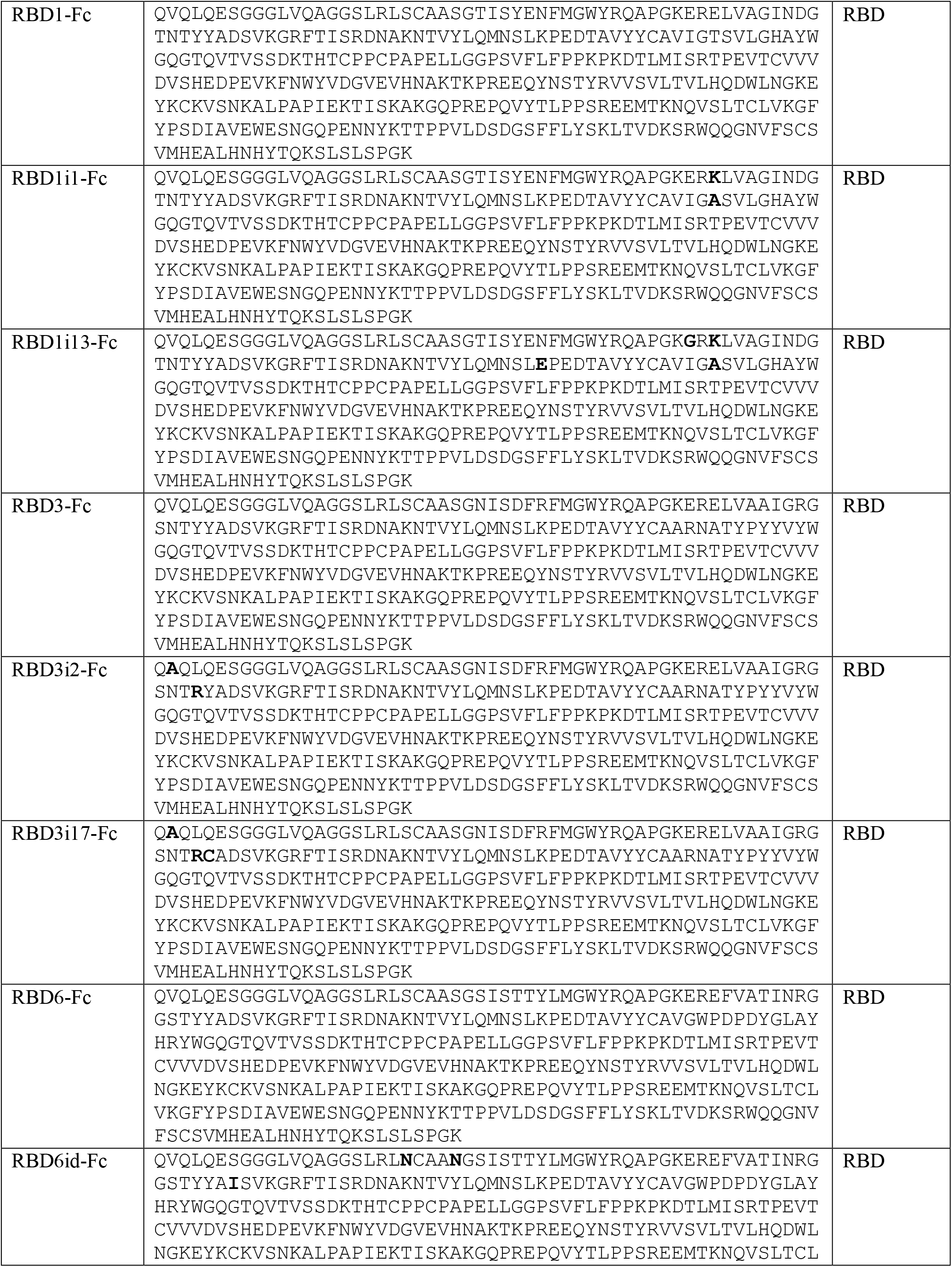
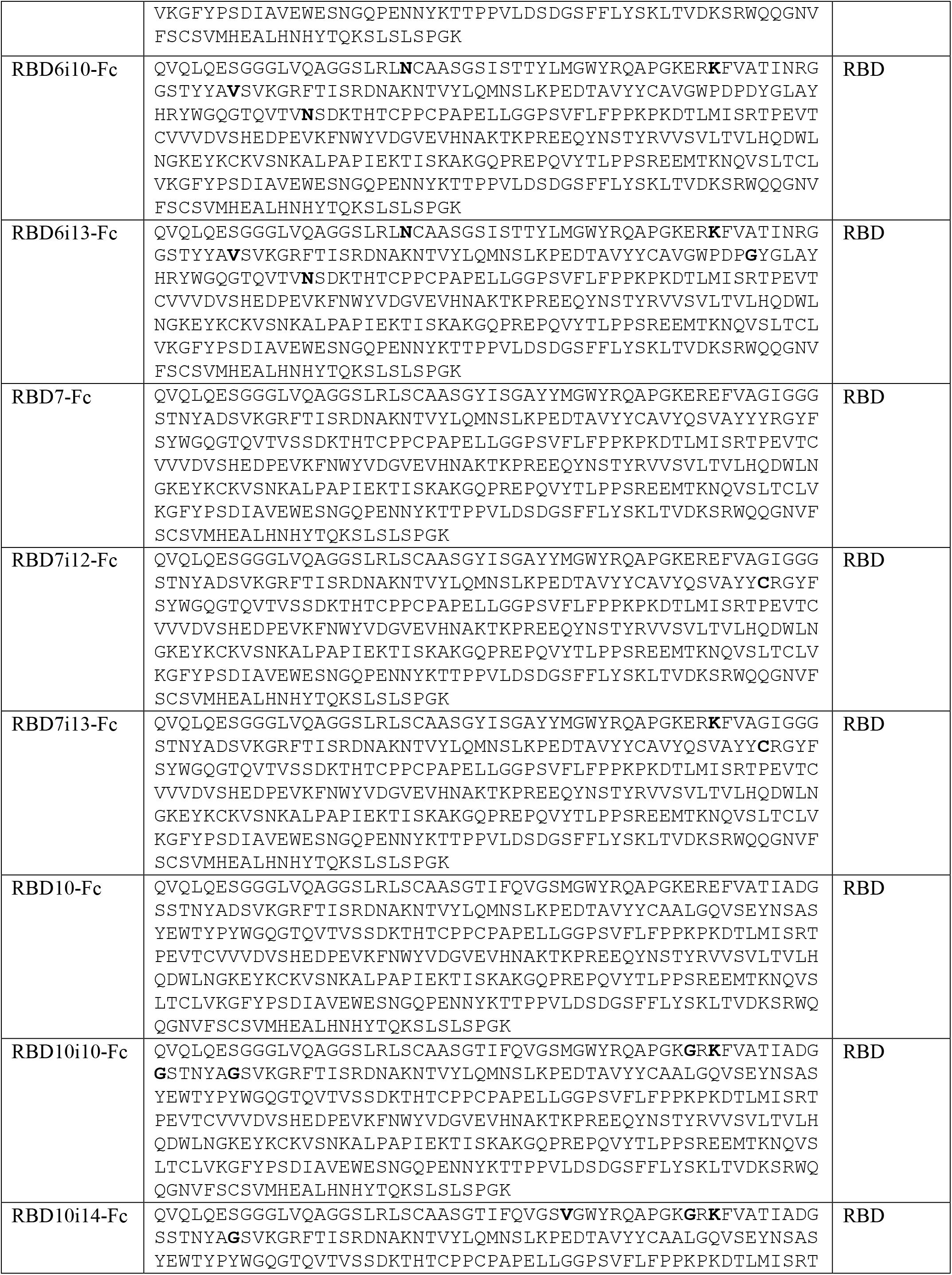
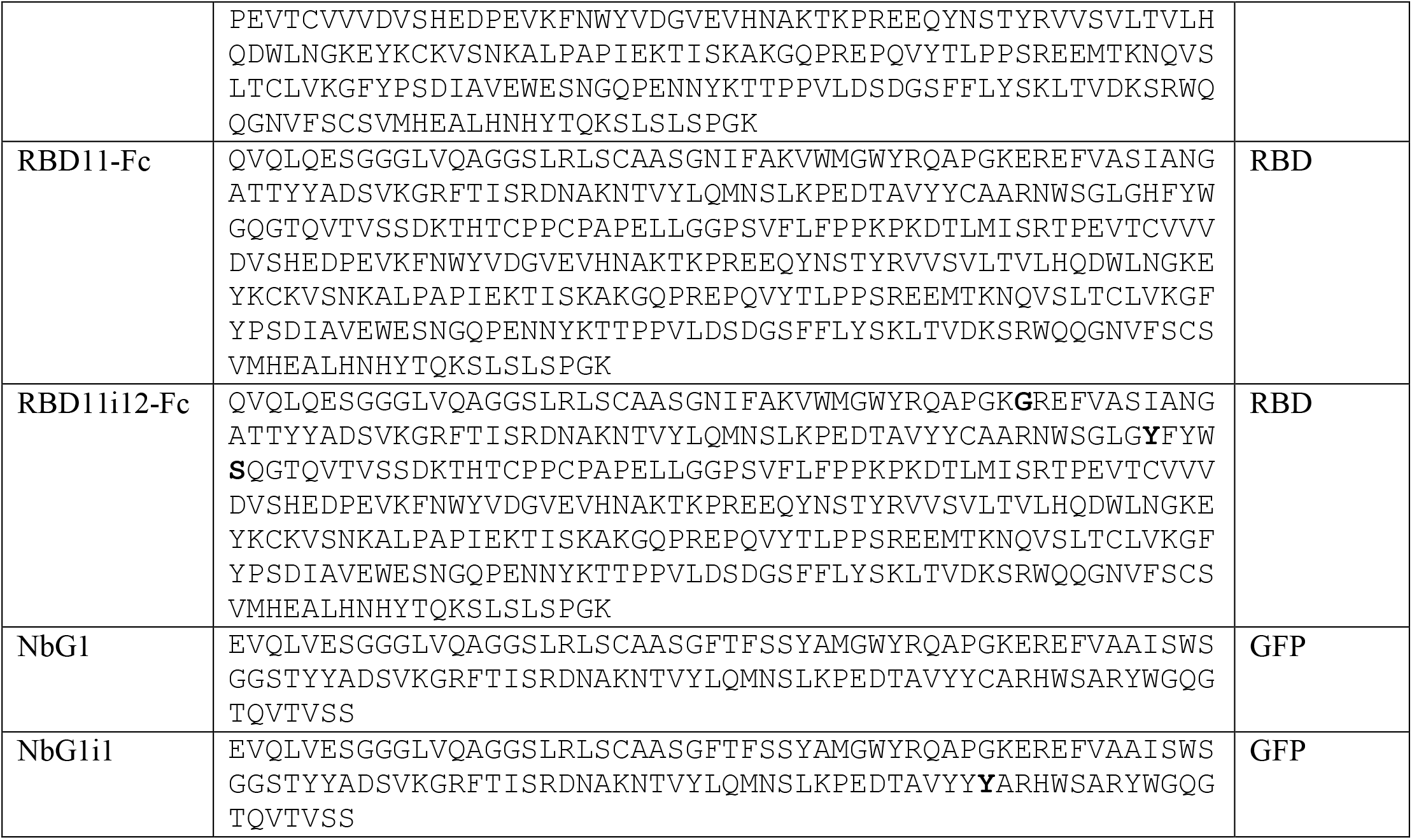
Sequences of all nanobodies described in this study.

**Table S3.** List of key plasmids used in this study

| **Name** | **Details** | **Source** |
| --- | --- | --- |
| pGKL1-short-M17 | Integration plasmid for replacing wild-type p1's immunity genes with MET17 in order to generate the landing pad p1 for transfer into other strains and ease of nanobody integration onto p1. | This study |
| pAR-633-Leu2 | A yeast CEN/ARS nuclear plasmid that expresses the error prone TP-DNAP1-4-2 with a Leu2 marker for selection. | This study |
| pAW24 | Integration plasmid for encoding nanobodies onto p1 in the AHEAD 1.0 format. In the parent plasmid, the nanobody AT110 is encoded in the integration cassette in the plasmid, but this nanobody can be replaced with any nanobody of interest through subcloning. The linear integration cassette generated from this plasmid by ScaI digestion integrates into p1 or landing pad p1 while deleting immunity genes (in the case of p1) and the TP-DNAP1 (in the case of p1 and landing pad p1). | This study |
| pAW240 | Integration plasmid for encoding nanobodies onto landing pad p1 in the AHEAD 2.0 format. In the parent plasmid, the nanobody AT110 is encoded in the integration cassette in the plasmid, but this nanobody can be replaced with any nanobody of interest through subcloning. The linear integration cassette generated from this plasmid by ScaI digestion integrates into landing pad p1 while deleting the TP-DNAP1 on the landing pad p1. | This study |
| pFUSE-hIgG1-Fc | Nanobody-Fc fusion expression plasmid. All RBD nanobodies characterized by SPR and neutralization assays were subcloned into this plasmid, expressed, and purified. | InVivoGen |
| FDP-P10B2-A75-RZ-URA3 | Source for polyA-RZ to augment nanobody expression. | Chang Liu, Addgene #130874 |
| pYDS649 | Plasmid for the yeast surface display of a naïve nanobody library. | Andrew Kruse, doi: 10.1038/s41594-018-0028-6 |
| pCAGGS-SARS-CoV-2 | Plasmid used to generate lentivirus pseudotypes. | This study |
| psPAX2 | Lentiviral packaging vector containing HIV Gag, Pol, Rev, and Tat. | Didier Trono, Addgene #12260 |
| pLenti-EF1a-Backbone | pLenti transfer vector containing GFP. | Feng Zhang, Addgene #27963 |
| pHLsec-SARS-CoV-2-RBD | Expression plasmid for recombinant SARS-CoV-2 RBD production. | This study |
| pVRC8400-RBD | Expression plasmid for recombinant RBD production. | Aaron Schmidt: doi: 10.1126/sciimmunol.abe0367 |
| pVRC8400-hACE2 | Expression plasmid for recombinant human ACE2 production. | This study |
| p253-Nb | Yeast nuclear plasmid for the galactose-inducible surface display of nanobodies in EBY100. Nanobodies were | This study |
|  | cloned into this plasmid in fresh EBY100 strains for on-yeast EC<sub>50</sub> measurements of individual nanobody clones. |  |
| pAW258 | Yeast nuclear plasmid encoding AT110 nanobody with an app8 secretory leader controlled by a SAC6 promoter for surface display of AT110 in EBY100. This was used to prepare an error-prone library of mutated app8 leader sequences for selection of improved leader sequences. | This study |
| pAW240-Nb.b201 | P1 integration plasmid for nanobody Nb.b201 (HSA binder). | This study |
| pAW240-Lag42 | P1 integration plasmid for nanobody Lag42 (GFP binder). | This study |
| pAW240-NbCM | Scaffold plasmid for CDR3 library cloning and further integration into yAW301. |  |
| pAW240-RBD1 | P1 integration plasmid for nanobody RBD1 | This study |
| pAW240-RBD3 | P1 integration plasmid for nanobody RBD3 | This study |
| pAW240-RBD6 | P1 integration plasmid for nanobody RBD6 | This study |
| pAW240-RBD7 | P1 integration plasmid for nanobody RBD7 | This study |
| pAW240-RBD8 | P1 integration plasmid for nanobody RBD8 | This study |
| pAW240-RBD9 | P1 integration plasmid for nanobody RBD9 | This study |
| pAW240-RBD10 | P1 integration plasmid for nanobody RBD10 | This study |
| pAW240-RBD11 | P1 integration plasmid for nanobody RBD11 | This study |
| pGLO-HIS6 | Bacterial expression vector for expression and purification of GFP with a C-terminal hexa-histidine tag. | This study |

**Table S4.**
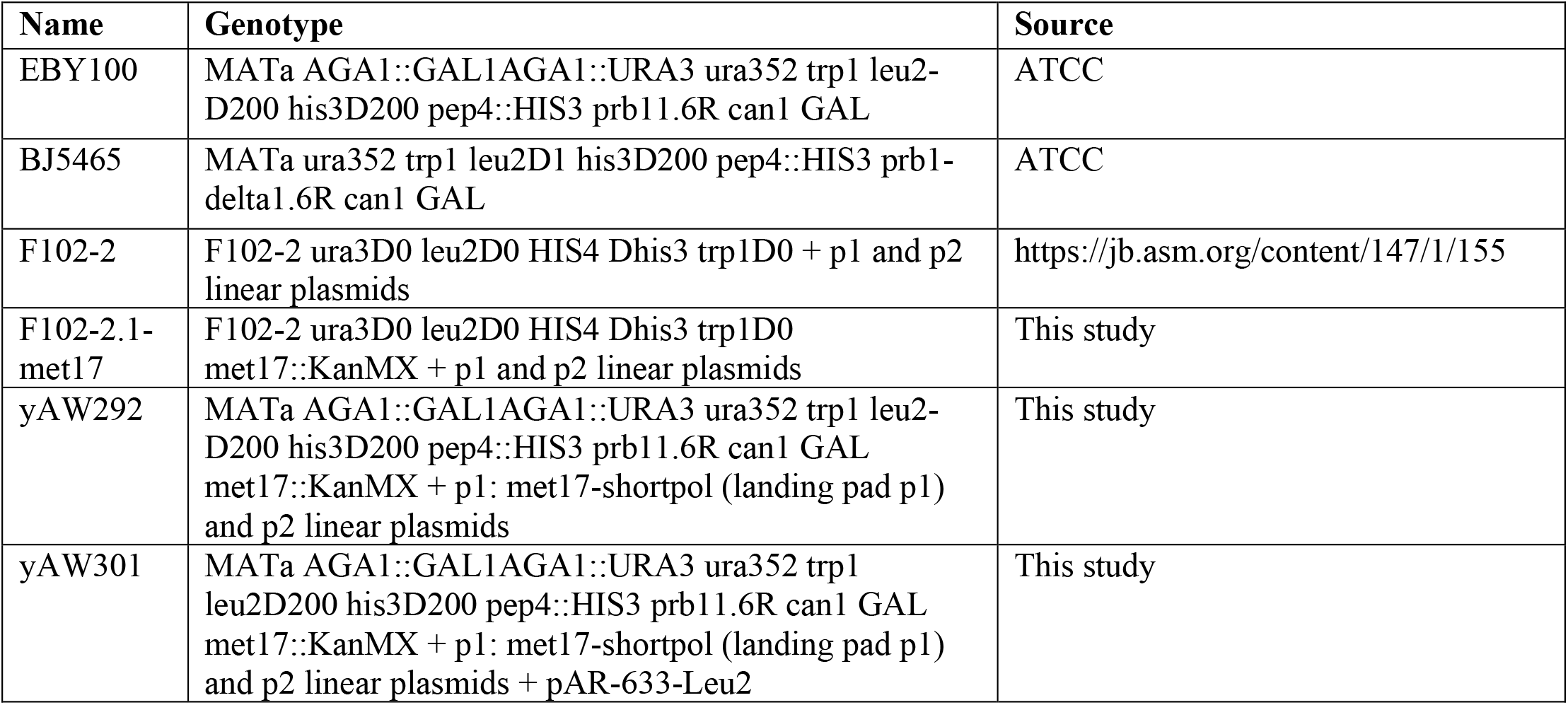
List of key yeast strains used in this study.

**Table S5.**
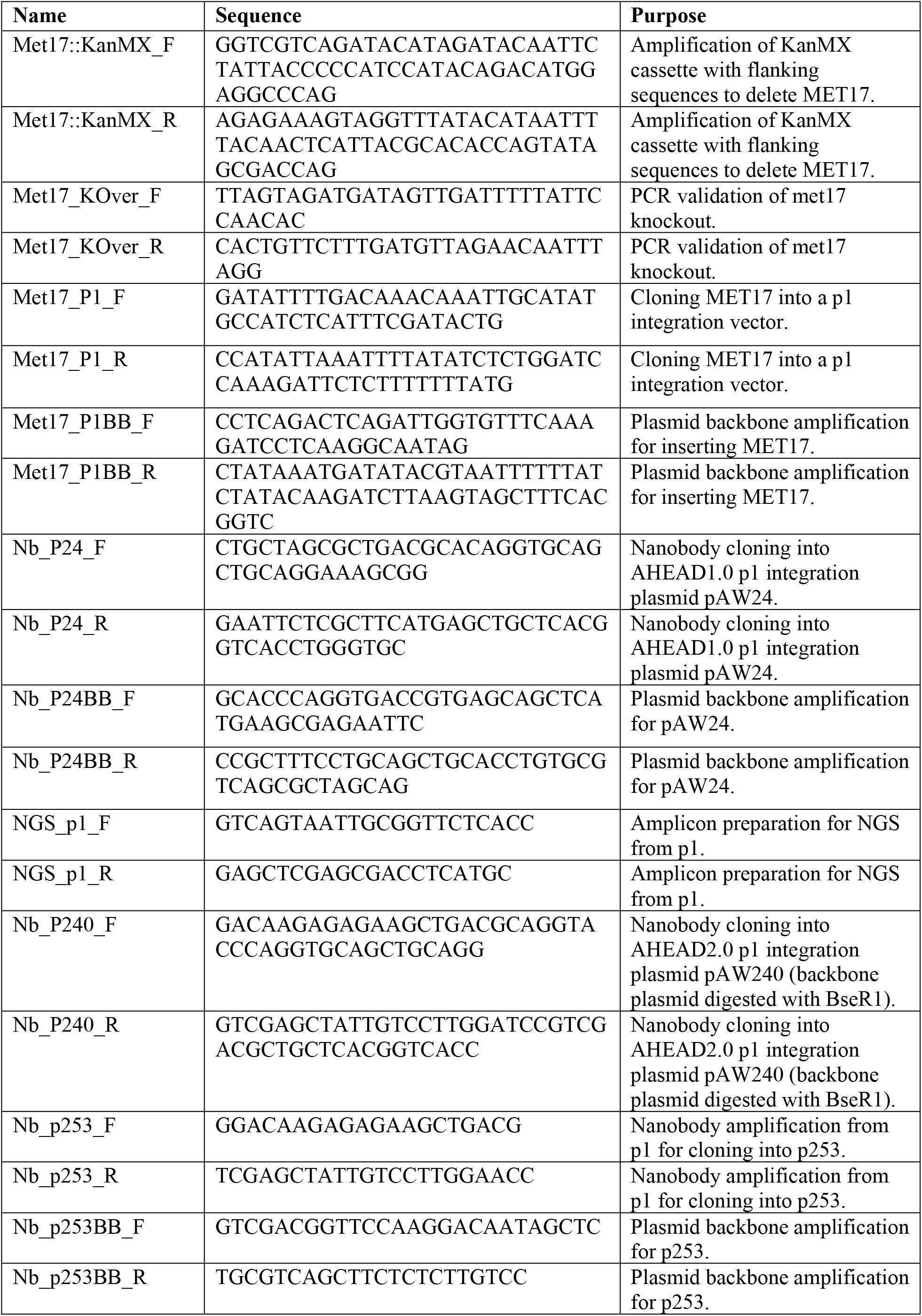
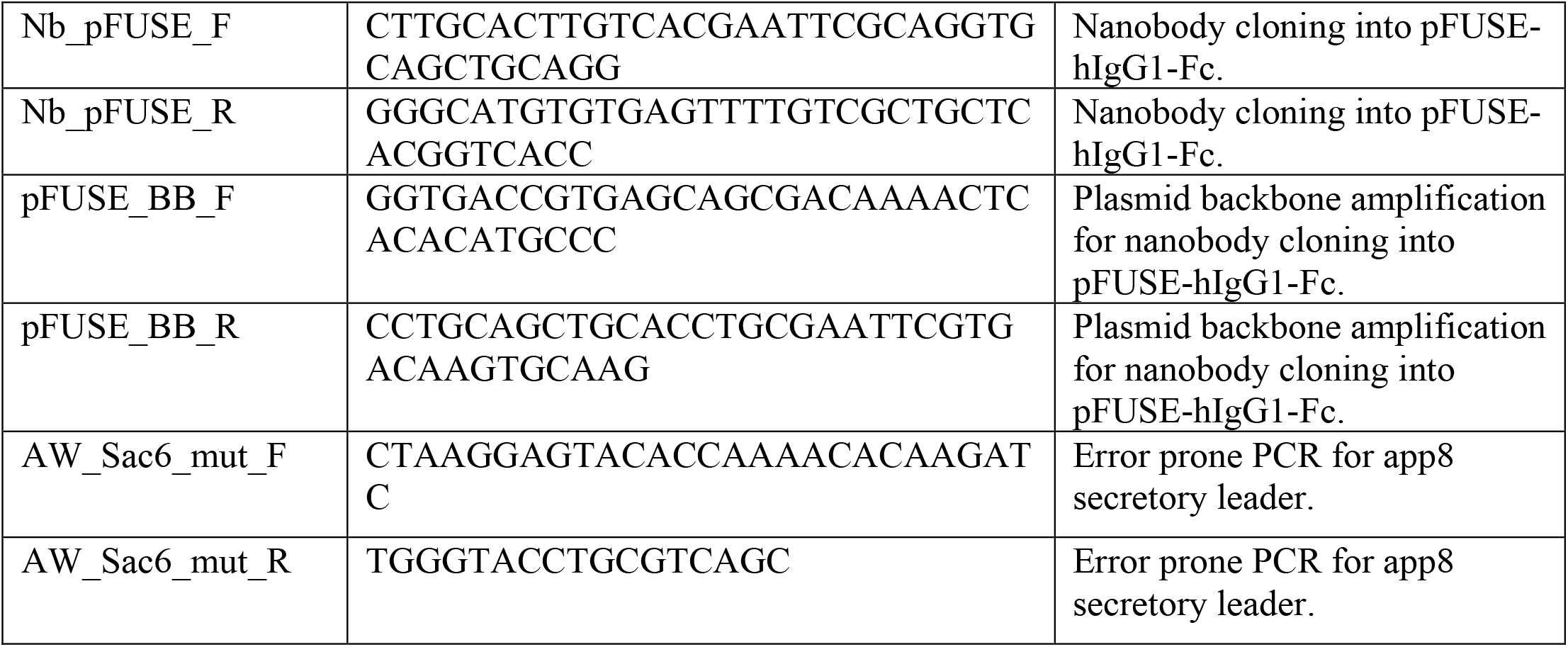
List of key primers used in this study.

